# Interconnecting ADC Structure with Tumor Cell Biology with Multimodal Learning

**DOI:** 10.64898/2026.05.22.727320

**Authors:** Hazem Mslati, Malcolm G. Wilson, Melika Naeinipour, Gaël Coulombe, Mehdi Ezzine, Tiana Yuen, Omar Abdal Bari, Hardeep Singh, Roger Tam, Joey G. Sheff, Francesco Gentile, Jeffrey V. Leyton

## Abstract

Antibody-drug conjugates (ADCs) represent a significant advancement in cancer therapy, yet their development remains constrained by high attrition rates driven by an incomplete understanding of how ADC chemical design interconnects with tumor biology. Compounding this challenge, the field has converged on a narrow set of redundant structural components, and current linker-payload systems that do not share a single mechanism of action. Existing drug response prediction frameworks cannot resolve this multidimensional complexity, relying predominantly on genomic inputs while protein-level biology is challenging to integrate. To address this, we developed a multimodal machine learning platform interconnecting ADC structural parameters with tumor cell biology across thousands of curated structure-activity datapoints, including multi-omics profiles from 1,479 human tumor cell lines and protein-level inputs from a unique model (GENCEP) that derives complete proteomic signatures. The model was validated through blinded retrospective evaluation and, critically, large coverage prospective prediction of cytotoxicity across 159 ADC-cell line combinations spanning five antigens, four mechanistically distinct linker-payload systems, and eight tumor types, most with no prior published associated ADC data. Performance surpassed industry benchmarks established for small molecule therapeutic modalities, demonstrating that protein-informed multimodal integrated framework is effective at capturing cytotoxic determinants at scale.

## Main text

Designing effective antibody-drug conjugates (ADCs) requires solving a multivariable problem that current strategies cannot scale to address. Although ADCs have transformed targeted cancer therapy by combining monoclonal antibodies with cytotoxic payloads through specialized linkers, their development remains slow, costly, and marked by high attrition (*1, 2*). Each ADC integrates interdependent design variables, including antibody antigen specificity, linker chemistry, payload class, at optimized drug-antibody ratios (DARs), whose therapeutic performance depends on tumor-specific processes such as antigen expression density, internalization kinetics, intracellular transport, lysosomal degradation efficiency, and cellular sensitivity to trigger apoptosis. These biological determinants vary widely across cancers, disease states, and normal cell counterparts, and although, linkers have advanced to exploit cellular underpinnings (*3*), the integration of cellular variables with ADC structural parameters tailored for specific cancer types has not been achieved, as each has been studied largely in isolation (*4–10*). A further dimension of this complexity is that ADC linker-payload systems do not share a single mechanism of action (MOA). While canonical ADCs depend on antigen-mediated internalization and intracellular payload liberation, certain linker chemistries undergo rapid hydrolysis under physiological conditions, where payload release is not strictly dependent on receptor engagement and function as prodrug-like delivery systems with distinct biological determinants of tumor sensitivity (*11–17*).

Paradoxically, despite this complexity, ADC development explores only a narrow fraction of the potential design space (*2*). The field exhibits substantial component redundancy, with repeated reliance on a limited set of linker-payload chemistries and canonical target antigens (*18*). This constrained exploration reflects both technical feasibility and economic pressure where comprehensive combinatorial testing across diverse ADC designs and tumor contexts is impractical. As a result, most programs proceed through *in vitro* screening, where single-to-few-component changes are evaluated for their influence on cytotoxicity through limited cell line testing (*19–21*). While this empirical approach has yielded successful therapeutics, it cannot systematically and comprehensively interrogate how ADC chemistry interacts with tumor biology across different tumors. This scaling challenge contributes to costly failures, delayed timelines, missed therapeutic opportunities, and unreliable therapeutic outcomes (*18, 22, 23*). Even after approval, ADCs have been withdrawn due to efficacy ineffectiveness (*24, 25*) or unanticipated severe toxicities (*26–28*), both traceable to the limited interconnection between ADC design and important biological underpinnings.

Artificial intelligence, specifically machine learning (ML) models have been proposed as potential solutions to optimize development across diverse therapeutic modalities (*29*), including ADCs (*30*). Most frameworks integrate tumor genomic and transcriptomic profiles with molecular drug descriptors, yet in-depth protein-level biology remains absent despite drugs acting at the protein level. Although biology-focused datasets are emerging (*31–33*), model generalization is typically demonstrated using known historical data benchmarks where predictions are assessed retrospectively (*4, 34–37*). More rigorous approaches validate computationally against independent unseen datasets (*38*), yet these remain bounded by available data. The central unmet need is therefore more meaningful biological contextualization of model inputs, coupled with a validation standard that reflects real-world predictive utility. This limitation is particularly relevant for oncology drug development due to the widely diverse mechanisms driving cancer (*39*). Prospective wet-lab validation represents one of the most stringent benchmarks for evaluating ML-based drug design strategies prior to translational commitment, yet such studies remain rare due to the absence of standardized experimental frameworks and the structural and mechanistic complexity of biologics, such as ADCs.

To address this, we developed the ADC Design Platform, a ML framework that comprehensively interconnects ADC structural design with tumor cell biology to enable rational, biology-informed screening and optimization (Fig. 1, Figs. S1-S2). The platform is built on a binary classifier, predicting whether an input ADC design is likely to be active or inactive for a defined tumor cell line at a clinically relevant cytotoxic potency threshold. The platform is composed of three separate components we created.

**Fig. 1.**
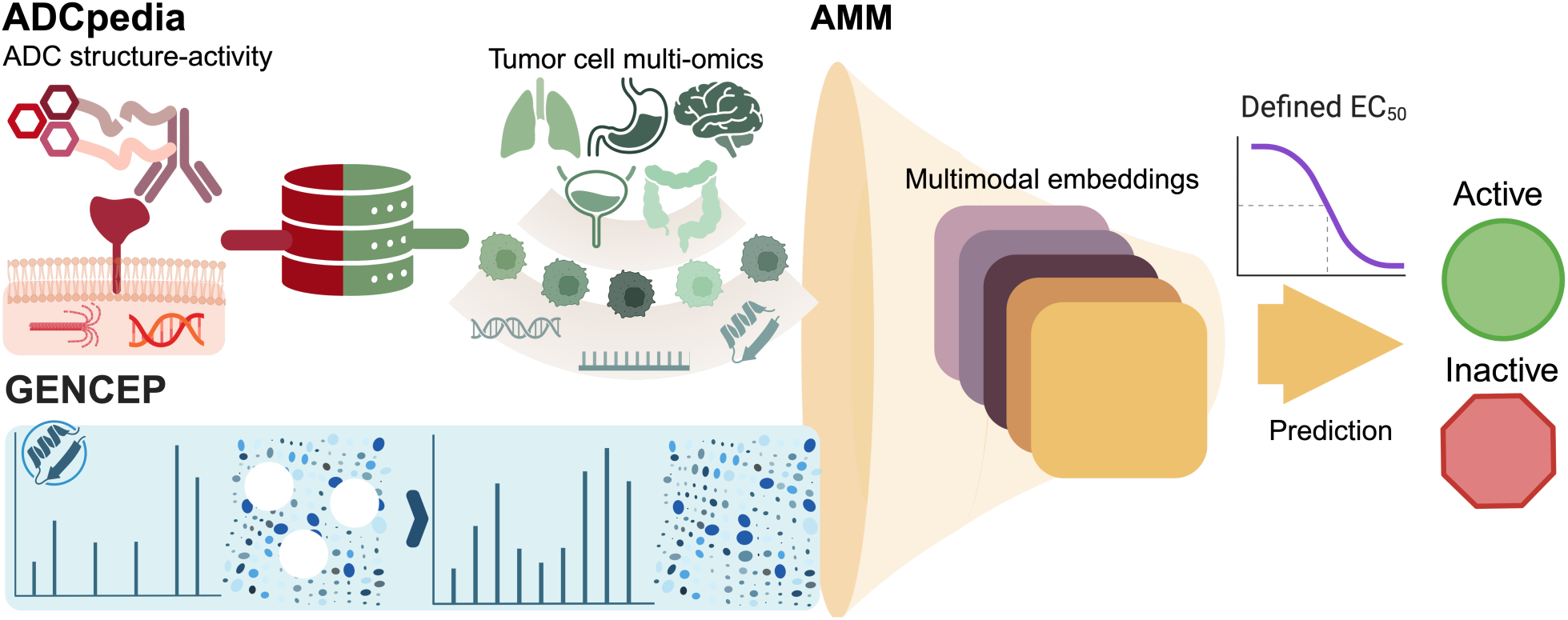
The multimodal framework of the ADC design platform. The platform enables the prediction of ADC activities across human tumor cell lines and is comprised of three components. ADCpedia is a curated database that includes records on ADC structures and cytotoxic EC_50_ activity values. It also contains 1,479 human tumor cell lines and their corresponding multi-omics data. The GENe Cell Expression Prediction (GENCEP) model addresses transcriptomics to protein expression gaps and provides biological information-rich, pretrained cell line embeddings. The ADC MultiModal (AMM) model is trained with this information at a 10 nM EC_50_ threshold and generates activity probabilities for an ADC design-tumor cell line combination.

First, ADCpedia is a highly curated database from analyzing nearly 25 years of literature on ADC structural-activity relationships. It is also coupled to a large multi-omics dataset from 1,479 human tumor cell lines (including 12 non-cancer cell lines), representing 98 tumor types. As a result, ADCpedia is a repository that converted fragmented historical ADC knowledge into an actionable design space, where users can cross-reference prior linker-payload classes, tumor contexts, and observed cytotoxicity performance to define design options and screening strategies with increased precision.

Second, GENCEP (GENe Cell Expression Prediction) addresses a fundamental bottleneck in biology-informed predictive modeling, specifically the pervasive incompleteness of proteomic datasets. This is critical as protein abundance, and not mRNA expression, is the primary driver of cellular processes governing ADC MOA and cellular response. GENCEP imputes missing proteomic intensities across the cell lines in ADCpedia and can provide protein intensity profiles important for predictive modeling and targeted profiling across the collection of tumor types in ADCpedia.

Third, the AMM (ADC Multimodal Model) is a deep learning (DL) model that integrates ADCpedia and GENCEP data to process the interconnections between ADC structure and tumor cell biology to determine cytotoxic response, classifying each ADC-tumor cell line context as active or inactive at 10 nM (*i.e*., ≤10 nM = active).

We demonstrate that the interconnection between specific ADC physicochemical parameters and mechanistically relevant protein expressions within an integrated tumor cell biological framework is an effective approach that successfully identified active and inactive ADC-cell line combinations. Performance was significant in a blinded retrospective test set and exceeded currently known industry wet-lab benchmarks for therapeutic drug discovery when the platform was prospectively validated across 159 ADC-cell line combinations spanning five major target antigens (HER2, Trop2, Nectin-4, CD19, and CD22) and four linker-payload systems (vc-MMAE, CL2A-SN38, vc-SN38, and GGFG-DXd) representing mechanistically distinct cytotoxic modalities, from stable protease-cleavable systems, to those more susceptible to hydrolysis. The nine cell lines that were tested represented eight tumor types (breast, colorectal, ovarian, acute lymphoblastic leukemia, acute myeloid leukemia, cervical, mesothelioma, and bladder), and seven of them had no prior published ADC data, directly challenging the platform in previously unchartered biological territory. Predictions were deliberately stress-tested across the full spectrum of biological contexts – from predicted sensitive to mixed response to resistant tumor models, the latter is where rational design tools add the most value. The platform is accessible through an online interface, making this biological landscape and predictive capability available to the global ADC community, and providing the field with a comprehensive decision framework for early ADC development.

## RESULTS

### A multimodal framework for ADC activity prediction

Analysis of the historical ADC landscape revealed that 46% of all reported ADCs had no associated biological testing data, and a further 28.4% had only cell line or target identification data, with only 5.5% reaching clinical trial pipelines or approval (Fig. 2A). Structural and biological diversity analysis further revealed widespread redundancy for the field, with over 60% of reported ADCs relying on redundant antibody, linker, payload, and antigen combinations (Fig. 2B). Heavy reuse of ADC components reflects the economic reality of ADC development, where a typical ADC requires on average 10 years and an investment of USD 1-2 billion from design to FDA approval for a single indication (*40*). While this strategy reflects economic constraints, it may also contribute to high ADC attrition (*18*), highlighting the tension between cost containment and comprehensive optimization. This pressure is amplified when high-stakes asset *go/no go* decisions must be made early in preclinical development, where data remains incomplete and biological context is poorly resolved. With over 370 ADCs now in clinical trials (*41*), this expanding pipeline highlights not only the momentum of the field, but also the vast number of potentially informative design-biology combinations that remain untested or underexplored.

**Fig. 2.**
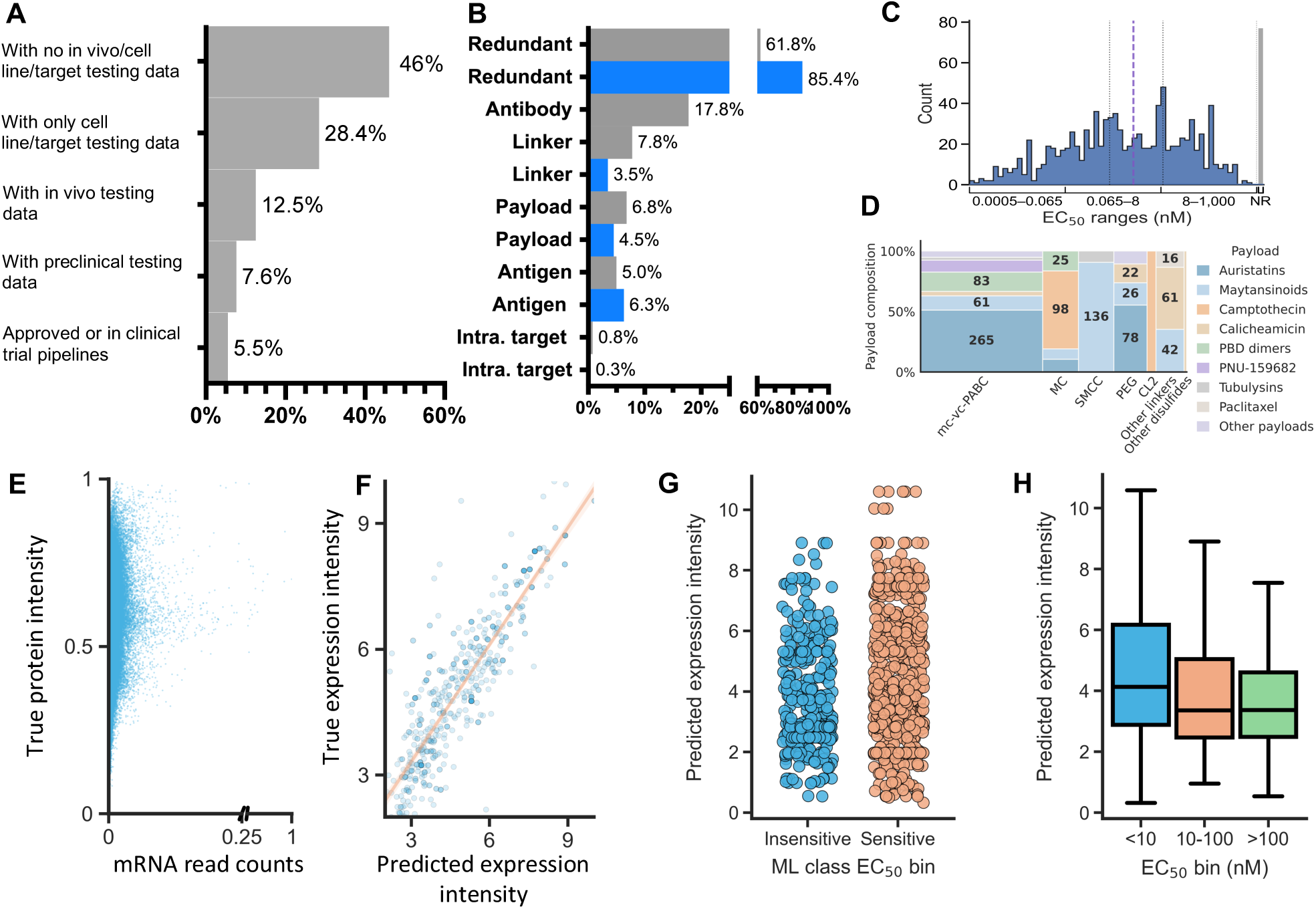
The ADC development landscape and the GENCEP model. (**A**) Proportion of historical ADC designs and development advancement stages. (**B**) Historical published (gray) and records in ADCpedia (blue) of percentage of unique ADC components and cell surface and intracellular targets. (**C**) Bimodal EC_50_ distribution for all published ADCs with activity records since 2001. Gray bar = >1000 nM, considered not reached (NR) and not incorporated into determining average for ADC cytotoxicity. (**D**) The distribution of linker-payload systems deployed in ADCpedia. (**E**) Correlation between protein expression intensities and mRNA counts. (**F**) Correlation between GENCEP-predicted protein intensities of ADC target antigens and ground-truth values. (**G**) Point-biserial correlation between EC_50_ values represented by binary values (insensitive >10 nM and sensitive ≤10 nM) and GENCEP-predicted antigen intensities. (**H**) Box plots of GENCEP-predicted antigen intensities stratified by high, mid, and low expression and their relationships with ADC EC_50_ values.

The scale and redundancy of the ADC landscape also reveal a more practical problem for using a computational approach to aide ADC development. Existing frameworks have relied on datasets assembled without requirements for complete structural specifications or paired quantitative cytotoxicity data and have not incorporated relevant biology inputs (*2, 42*). Broad data aggregation without stringent curation leaves gaps that are detrimental to model training and obscure the design-activity relationships needed for meaningful interrogation.

To address this, ADCpedia was constructed using a curation strategy distinct from previous approaches. To build a training foundation appropriate for ML, the curation began from preclinically and clinically validated ADCs catalogued in the ADC DrugMap (*43*). Records were filtered to include only canonical ADC architectures with complete structural specifications and paired to quantitative cytotoxicity data generated under standard assay conditions (Materials and Methods), which enabled a targeted literature search from January 1, 2000, to April 15, 2024. This process yielded 1,167 curated datapoints from 151 sources and captured 200 structural identity and physicochemical features for each ADC component, including linker-payload compositions, conjugation sites, and DAR (Materials and Methods). Each record was further annotated with the corresponding target antigen, tumor type, cell line name, and EC_50_ value. Critically, the dataset was coupled to a comprehensive multi-omics dataset from 1,479 human tumor cell lines (including 12 non-tumor cell lines) representing 98 cancer types ((*44*); Fig. S1). ADC structural and biological omics features were encoded as embeddings connecting ADC design parameters with tumor cell biology, forming the input representation for model training (Materials and Methods).

Analysis of EC_50_ values across ADCpedia revealed a bimodal activity distribution, with two distinguishable high (∼0.2 nM) and low (∼35 nM) activity populations resulting in a natural separation at approximately 10 nM (Fig. 2C). This potency divergence reflects a historical shift in ADC design philosophy: from early constructs bearing traditional chemotherapeutic payloads toward modern ultrapotent payload chemistries that emerged in the 2000’s, when the field moved away from micromolar-range payloads to payloads achieving cytotoxicity in the picomolar-to-low-nanomolar range (Fig. S3A). Strikingly, this wide potency range coexists with a remarkably narrow structural design space – auristatins, maytansinoids, and camptothecin derivatives paired with only a handful of linker chemistries account for most of all ADC-cell line combinations in nearly 25 years of ADC development (Fig. 2D). Closer inspection within the modern high-potency population reveals that EC_50_ values vary more than 300-fold, spanning from approximately 0.1 nM to over 30 nM (Fig. S3B). This wide distribution within this ‘potent’ ADC era highlights a key unresolved question of what constitutes an effective ADC design for a given antigen-tumor cell line combination and is further confounded by diverse cytotoxicity assay conditions, which was not only challenging for curation (Materials and Methods) but also for optimizing real-world validation studies (detailed further below in prospective validation result section).

Contextualizing the relationships between EC_50_ values and individual ADC design components using ADCpedia provides deeper granularity for this variability. For example, stratified analysis by payload class revealed dramatic EC_50_ divergences across tumor types within the three main payload categories (microtubule inhibitors, DNA damagers, and topoisomerase I inhibitors), with both high- and low-potency outcomes depending on the tumor type context (Fig. S4). Linker chemistry also revealed a wide variability with cleavable linker-based ADCs (Fig. S5A). In contrast, most non-cleavable linker-based ADCs were potent (Fig. S5B). DAR values spanning 0.8 to 9.0 similarly failed to reveal consistent potency trends across tumor types (Fig. S6). Taken across all three structural variables, widespread variability is the dominant pattern, where no single parameter, nor any combination of payload class, linker chemistry, and DAR reliably associated cytotoxic potency across tumor types. Structural ADC parameters alone are therefore insufficient to explain cytotoxic outcome and, perhaps, tumor cell biology is the missing determinant.

### Closing the transcriptomics-to-proteomics gap with GENCEP

Protein abundance is critical for the mechanistic process by which ADCs perform (*5, 23*). Although cellular proteomics quantifies protein abundance, a key barrier to biology-informed ML modeling is the incompleteness of proteomic data. The large gaps in protein intensities are attributable to degradation, post-translational modifications, and inadequate capture by current technologies (*45*). Additionally, protein intensities and mRNA levels are poorly correlated and cannot be scaled from transcript counts (*44, 45*). Consistent with this, protein intensities across the cell lines in ADCpedia showed poor correlation with mRNA transcript levels (Fig. 2E), with approximately 91% of proteomic intensity data absent relative to transcriptomic counterparts, with pronounced gaps for ADC-relevant target antigens (Fig. S7A).

GENCEP was developed to address this bottleneck to enable proteome-wide prediction of missing protein intensities, validated against ground truth experimental values prior to integration into ADCpedia (Materials and Methods; Fig. S7B). On held-out test data, GENCEP predicted protein intensities with R^2^ = 0.87 (Fig. S7C), and R^2^ = 0.75 specifically for ADC target antigens (Fig. 2E), substantially outperforming a zero-rule regression model (R^2^ = -8 x 10^-6^). Expanding past antigens, GENCEP also provides universal standardization of protein abundance levels across cell lines, enabling quantitative integration of any protein across the cell lines in ADCpedia.

With GENCEP’s ability to predict antigen expression across all tumor cell lines in ADCpedia, we aimed to further clarify whether antigen intensity alone could stratify ADC cytotoxic potency based on a 0–11-unit value metric pegged to the model antigen HER2, stratified by high (>9), mid (6–9), and low (<6) (Fig. S8 and S9A). Surprisingly, a poor point-biserial correlation (r = 0.145) between predicted intensities and binary active (≤10 nM) and inactive (>10 nM) classification confirmed no statistically significant association with potency (Fig. 2G), with substantial EC_50_ overlap across the three intensity bins for all ADC antigens in ADCpedia (Fig. 2H). This disconnect was consistent for HER2 (Fig. S9A), as well as additional well-characterized target antigens, such as Trop2 (Fig. S9B and Fig. S10) and CD79b (Fig. S9C), where expression-activity relationships overlapped, despite distinct expression bins. Evaluation of additional antigens was not possible due to insufficient accompanying cytotoxicity data. Moreover, leveraging this large-scale omics pretraining, both transcriptomic embeddings and GENCEP-derived antigen intensity predictions could also be integrated into ADCpedia and served as biologically informed input features for the AMM, thus augmenting the informational content of the otherwise limited ADC-specific training dataset.

The analyses establish a consistent and troubling picture. Nearly three-quarters of all ADCs ever developed have never been biologically tested for activity and the structural design space is dominated by a narrow set of redundant components, and yet, within that constrained landscape, cytotoxic potency varies by orders of magnitude in ways that neither payload class, linker chemistry, nor DAR can resolve. Proteomics is the most direct biological readout of the protein machinery governing ADC MOA and cellular response yet, is profoundly incomplete across the cell lines needed for ML training, and mRNA transcript levels cannot substitute for it. GENCEP addressed this by accurately predicting protein intensities and creating a standardization across all cell lines in ADCpedia, making quantitative cross-referencing of antigen expression against cytotoxic activity possible for the first time. Yet even with expression in hand, no meaningful association between antigen intensity and ADC potency emerges, further supporting that antigen expression is necessary but not sufficient. Therefore, what the ADC design-activity landscape demands is an approach that simultaneously integrates the complexity of ADC chemical architecture with a deeper tumor cell biological context to accurately determine whether a specific combination is ultimately lethal or not. That is the problem the AMM was built to solve.

### Accurate prediction of ADC bioactivity with AMM

The AMM is a multimodal late fusion convolutional neural network designed to predict ADC *in-vitro* activity by integrating ADC structural parameters with tumor cell biological information from ADCpedia (Fig. S2 and Table S1). Physicochemical representations of payload-linker systems are included. Specific cell line-wide transcriptomic embeddings derived from GENCEP are also included, as well as antigen Evolutionary Scale Modeling (ESM), predicted protein intensities, and read count embeddings for target antigens. Additional biological embeddings included GENCEP-predicted intensities and read counts for 27 ADC MOA-relevant proteins spanning mechanisms involved in ADC intracellular processing, payload sensitivity, and adaptive resistance (*46–48*) (Fig. S2A, Material and Methods). The AMM was trained as a binary classifier at a 10 nM EC_50_ threshold, consistent with the bimodal activity distribution identified in ADCpedia (Fig. 2C), with the training set constructed to ensure structural diversity across training, validation and test partitions (Material and Methods).

On the validation and held-out test sets, AMM achieved receiver operating characteristic-area under the curve (ROC-AUC) scores of 0.85 and 0.84, respectively (Fig. 3A). Confusion matrix analysis confirmed a statistically significant classification performance (*p* = 0.0004; Fig. 3B; Table S2), a specificity of 0.833 exceeding sensitivity of 0.667, indicating the model was stronger at correctly identifying inactive ADC-cell line combinations than active ones across the test set. The positive predictive value (PPV) of 0.70 and negative predictive value (NPV) of 0.811 indicate that the model’s active and inactive predictions were correct most of the time. The likelihood ratio (LR) of 4.0 indicates that a positive AMM prediction on the internal test set was four times more likely to reflect true ADC activity than true inactivity, establishing a discriminative baseline.

**Fig. 3.**
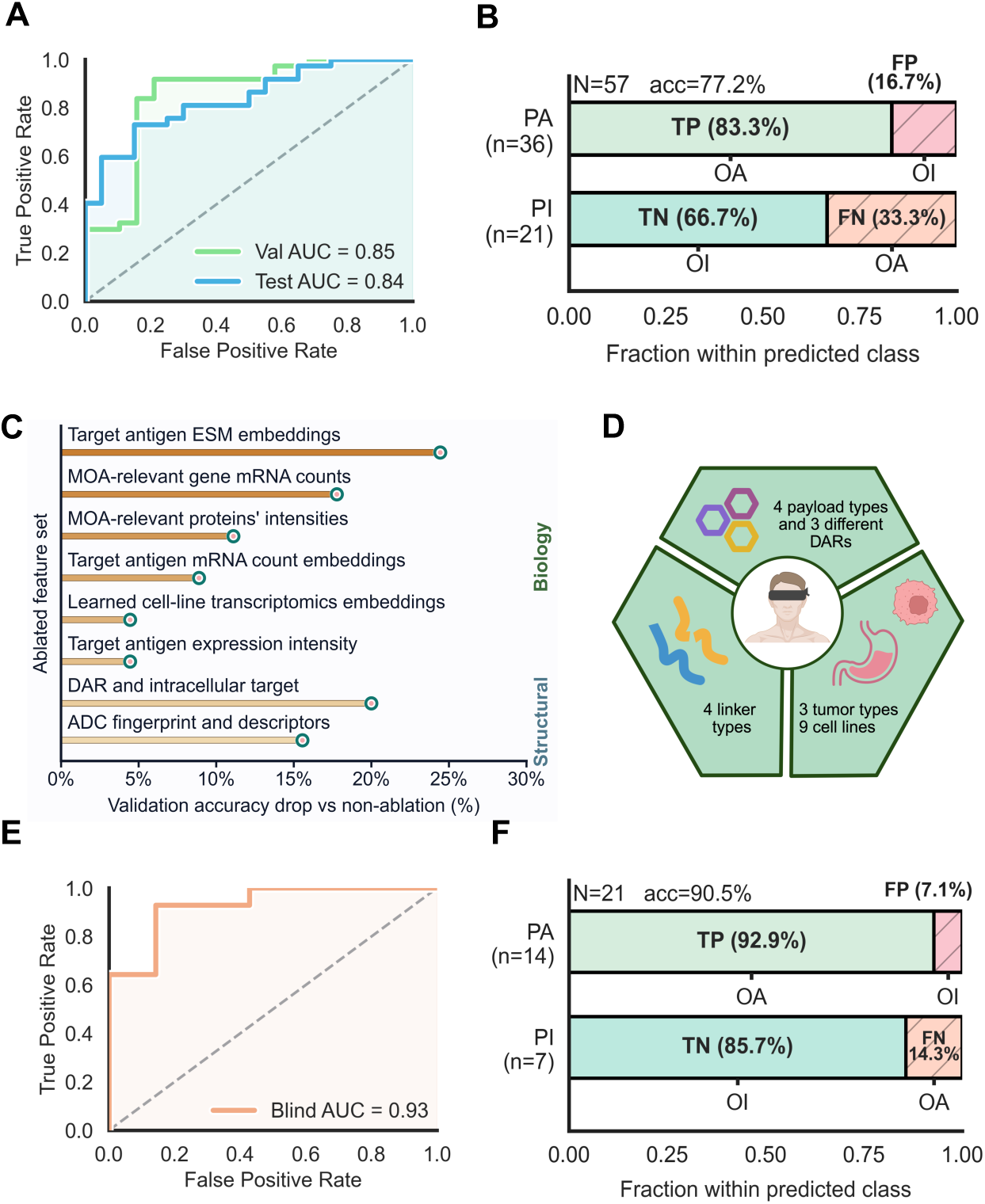
AMM performance on retrospective data. (**A**) ROC-AUC performances for the AMM on validation and hold-out (n = 57) test sets. (**B**) Confusion matrix for the held-out test set, where TP = true positive, FP = false positive, TN = true negative, FN = false negative, OA = observed active and OI = observed inactive. (**C**) The effect on AMM performance on the validation set when specific feature blocks were ablated. MOA = pooled 27 proteins relevant to ADC mode of action or implicated in chemotherapy resistance. (**D**) Schematic of the blinded retrospective evaluation. (**E**) ROC-AUC performance of AMM on novel ADC-cell line combinations (21) in the blind test set. (**F**) Confusion matrix for the retrospective evaluation on unseen combinations.

Ablation of individual feature blocks from both the biological and ADC structural sides confirmed that both factors are critical to AMM performance (Fig. 3C). Removing target antigen ESM embeddings produced the biggest accuracy drop at 25%. In contrast, ablating target antigen intensity only reduced performance by 5%. Pooled protein intensities relevant to ADC MOA and resistance, impaired AMM performance by 11%. The mRNA embeddings of target antigens and mRNA count of other proteins also contributed meaningfully to performance. ADC structural feature blocks, specifically DAR with intracellular targeting status or ADC fingerprint descriptors, also individually contributed 20% and 16% to AMM performance, respectively. This indicated that linker-payload properties also influenced performance distinct from the biological input elements. Taken together, these results demonstrate that AMM integrates biological and structural elements in a hierarchical fashion. Although direct ranking across feature blocks is not feasible given their distinct data types and organization with AMM’s architecture, both modalities were individually necessary. However, it appears as that protein-level information beyond target antigens is critical. Given the poor association between antigen expression and ADC potency (Fig. 2F and Fig. 2G), AMM’s predictive value emerged specifically from multimodal integration that incorporates a deeper tumor cell biological context beyond antigen expression.

To further evaluate AMM performance under realistic discovery conditions, the model was tested against an external unpublished dataset (Supplemental materials) where ADC activity was blinded to the authors prior to prediction (Fig. 3D and Table S3). This dataset comprised four structurally distinct HER2-targeted ADCs, T-DM1, T-DXd, disitamab-vc-MMAE, and trastuzumab-pAcF-Amberstatin 269, tested across nine tumor cell lines representing breast, esophageal, and gastric cancers, spanning three different DARs and four distinct linker-payload systems, for a total of 30 ADC-cell line combinations, 21 of which were unseen during training. On novel combinations, AMM achieved a ROC-AUC of 0.93 (Fig. 3E), with confusion matrix analysis revealing significant classification performance (*p* = 0.0009; Fig. 3F; Table S4). Specificity reached 0.857 and sensitivity 0.929. The PPV of 0.929 and NPV of 0.857 indicate that positive and negative predictions are each correct in nearly nine of ten cases. Notably, the LR of 6.5 indicates that a positive AMM prediction in the retrospective evaluation had a discriminative performance that exceeds the hold-out test, likely reflecting the model’s familiarity with HER2 biology relative to latter spanning multiple target antigens. Nevertheless, four investigated cell lines (UACC-812, EFM-192A, OE19, SNU-216) and one payload-linker system (trastuzumab-pAcF-Amberstatin 269) were entirely novel with respect to the training set, suggesting AMM’s potential to generalize to structurally and biologically novel ADC-tumor contexts.

### Broad and prospective validation of AMM across diverse tumor cell lines

Next, we prospectively validated the ADC Design Platform (Fig. 4A and workflow detailed in Supplemental materials). Five clinically relevant target antigens were selected spanning both hematologic and solid tumor indications: HER2, Trop2, Nectin-4, CD19, and CD22. Linker-payload systems were selected to represent the breadth of approved ADC chemistries and mechanistically distinct cytotoxic modalities, from classical protease-cleavable vc-MMAE to the more recent prodrug-like CL2A-SN38 and protease-cleavable GGFG-DXd, capturing ultratoxic and non-ultratoxic payload classes and different MOAs (*49*). DAR contributions to predictions were fixed at values reflecting relevant designs: CL2A-SN38 (DAR 7.1), vc-SN38 (DAR 3.5), GGFG-DXd (DAR 7.7), and vc-MMAE (DAR 2.2), with predictions additionally spanning DAR values classified as low (1–2), mid (3–5), and high (6–8) to confirm stability of classification outputs around these anchor DARs. Only subtle changes were observed, indicating linker-payload design and not DAR is critical.

**Fig. 4.**
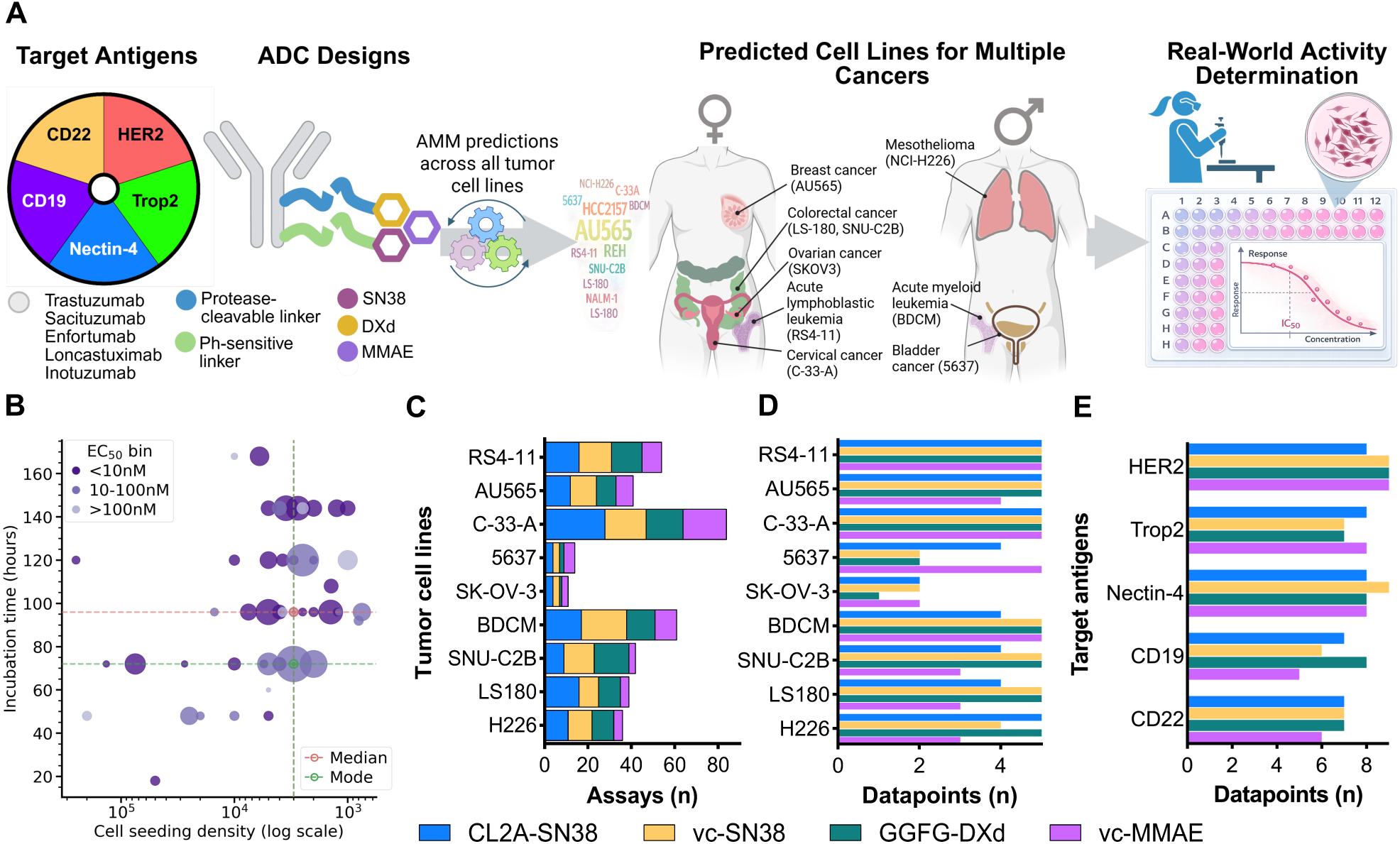
Prospective validation framework. (**A**) Evaluation workflow depicting the target antigens and ADC designs that were predicted against all cell lines in ADCpedia. Nine cell lines representing eight tumor types were selected based on availability. The ADCs were constructed, characterized and their cytotoxic EC_50_ values determined and compared to predictions. (**B**) Bubble plot capturing the relationships between cell density/well, ADC incubation time, and EC_50_ outcomes. Shade of purple is proportional to EC_50_ potency and number of assays from single-to-few-to-many is proportional to circle size. Red and green crosshairs indicate the mean and the most frequently observed (i.e., mode) associations, respectively. (**C**) Prospective assay coverage across cell lines and linker-payload systems (n = 377). (**D**) The number of averaged EC_50_ datapoints for each ADC-cell line combination, stratified by linker-payload system (n = 151). (**E**) Antigen coverage across linker-payload systems (n = 151).

Cell lines were identified from AMM prediction outputs, which provided activity probabilities across the full spectrum of tumor cell lines in ADCpedia for each of the above ADC designs. Cell lines were deliberately selected to span the full predicted probability range, importantly, cell lines with high predicted sensitivity, mixed probabilities, and low predicted sensitivity. In this way, the AMM could be stress-tested across a complete prediction range. Final selection was made on the basis of regional ATCC availability and standard media requirements for reproducibility (Table S5), also considering, where possible, cell lines that belong to different probability tiers could be evaluated in parallel with multiple ADCs. This yielded nine cell lines (six female, 3 male) across eight tumor types: breast cancer (AU565), colorectal cancer (LS-180, SNU-C2B), ovarian cancer (SKOV3), acute lymphoblastic leukemia (RS4-11), cervical cancer (C-33-A), mesothelioma (NCI-H226), acute myeloid leukemia (BDCM), and bladder cancer (5637), establishing a total of 180 unique target ADC design-cell line combinations that would enable a comprehensive evaluation of the interplay between ADC linker-payload design, target antigen context, and tumor cell biology. Critically, seven cell lines (5637, AU565, BDCM, C-33-A, SNU-C2B, LS-180, RS4-11), one payload-linker system (vc-SN38), and thirteen ADCs (antigen-linker-payload systems) were entirely novel not in the training pool, thus ensuring that AMM’s performance could not be attributed to overlap with well-characterized cell line and ADC systems, and that AMM predictions were generated in unchartered chemical and biological territory for the majority of this prospective evaluation. Moreover, prioritized ADC-cell line combinations were not specifically filtered to retain only pairs that were not seen during training; nevertheless, and strikingly, only a single ADC-cell line combination from the evaluated pairs was present in the training set (T-DXd tested on SKOV3).

The probability distributions across the selected combinations confirmed that the final cell line selection achieved broad and balanced tier representation. For instance, for the four linker-payload systems, the likely sensitive, mixed, and resistant prediction tiers were represented with no single system restricted to a narrow prediction range (Fig. S11). At the individual tumor cell line level, each selected cell line displayed a distinct probability profile for each linker-payload system, indicating AMM’s ability to resolve tumor cell line-specific differences rather than producing uniform predictions across linker-payload classes (Fig. S12). Viewed from the target antigen perspective, all five antigens were represented for all three prediction tiers, albeit CD19 and CD22 showed preferential clustering at sensitive- and resistant-tier predictions, respectively (Fig. S13). At the cell line and antigen intersection, there were common and distinct antigen-specific discrimination across tumor type cell lines (Fig. S14). For instance, hematologic malignancy cell lines RS4-11 and BDCM concentrated likely sensitive-tier predictions for CD19, while likely resistant-tier predictions were generated for solid tumor antigens. Interestingly, both RS4-11 and BDCM concentrated likely resistant-tier predictions for CD22, which is the target antigen for inotuzumab ozogamicin (an ADC design not evaluated in this study). Notably, the two colorectal cancer cell lines, LS-180 and SNU-C2B had similar patterns with almost all antigens predicted in the likely resistant-to-mixed-tiers, indicating these would be difficult to kill based on AMM predictions. In contrast, AU565 and SKOV3 had likely sensitive-tier predictions for HER2, Trop2, and Nectin-4, possibly, indicating breast and ovarian cancer biology may be more susceptible for these ADC designs.

Cumulatively, these distributions confirm that the prospective validation framework for cell line selection was broad and balanced for the three parameters central to ADC development – structural design, tumor cell context, and target antigen. Furthermore, this was achieved under practical real-world constraints rather than outcome optimization, greatly reducing bias for the platform’s evaluation.

A critical prerequisite to prospective validation was the standardization of cytotoxic assay conditions to ensure that platform performance could be attributed to AMM predictions rather than experimental variability. Therefore, assay parameters were benchmarked against the diversity of protocols represented in ADCpedia. A seeding density of 5,000 cells per well and a 72-h incubation period were established as two major conditions (Fig. 4B; Materials and Methods). Other conditions that were verified consisted of ADC construction quality, concentration range determination, cell culture procedures, specificity, EC_50_ calculation, and SN38 release kinetics (Materials and Methods and Figs. S15-S21).

Prospective cytotoxicity testing was performed on the nine cell lines against the linker-payload systems over a fixed six-month period under real-world laboratory conditions, where cell line availability, growth kinetics, and ADC construction/characterization timelines were not artificially controlled. This yielded 377 total cytotoxicity assays distributed across all ADC-cell line combinations, with all linker-payload systems represented for every cell line and the number of biological replicates per combination varying under practical experimental constraints (Fig. 4C and Table S6).

Each ADC-cell line combination contributed a single averaged EC_50_ datapoint regardless of the number of biological replicates performed, yielding a final 151 from the 180 possible unique datapoints within the prospective validation framework. Coverage was strong across most cell lines, with the majority approaching the maximum of five antigens per linker-payload system, demonstrating broad experimental representation across both structural and biological axes (Fig. 4D). Notably, 5637 and SKOV3 had partial coverage due to practical constraints during the experimental window, but all four linker-payload systems were still represented. The EC_50_ values with the corresponding prediction for each combination are provided in Table S6. The 151 datapoints are distributed across the five target antigens with balanced representation at 36 to 38 combinations per antigen for each linker-payload (Fig. 4E). There were an additional eight datapoints comprised of CL2A-SN38-based ADCs constructed at low, mid, or high DARs that had sufficient data for evaluation for a total of 159 data points. Together, the evaluated combinations and the resulting datapoints achieved balanced representation across all three dimensions, with the experimental coverage mirroring the predictive arm of the validation framework.

Across all 159 ADC-cell line combination data points, the AMM achieved a ROC-AUC of 0.73 (Fig. 5A), surpassing an industry benchmark range of 0.65-0.70 established for predictive models for therapeutic small molecules on single targets (*50*). Calibration analysis demonstrated a monotonic relationship between predicted probabilities and measured EC_50_ values, indicating that model confidence provided a meaningful estimate of cytotoxic outcome likelihood (Fig. 5B). This suggests that higher confidence predictions (cream tan bars) carry greater reliability and identifies a straightforward path to further precision improvement in future deployment.

**Fig. 5.**
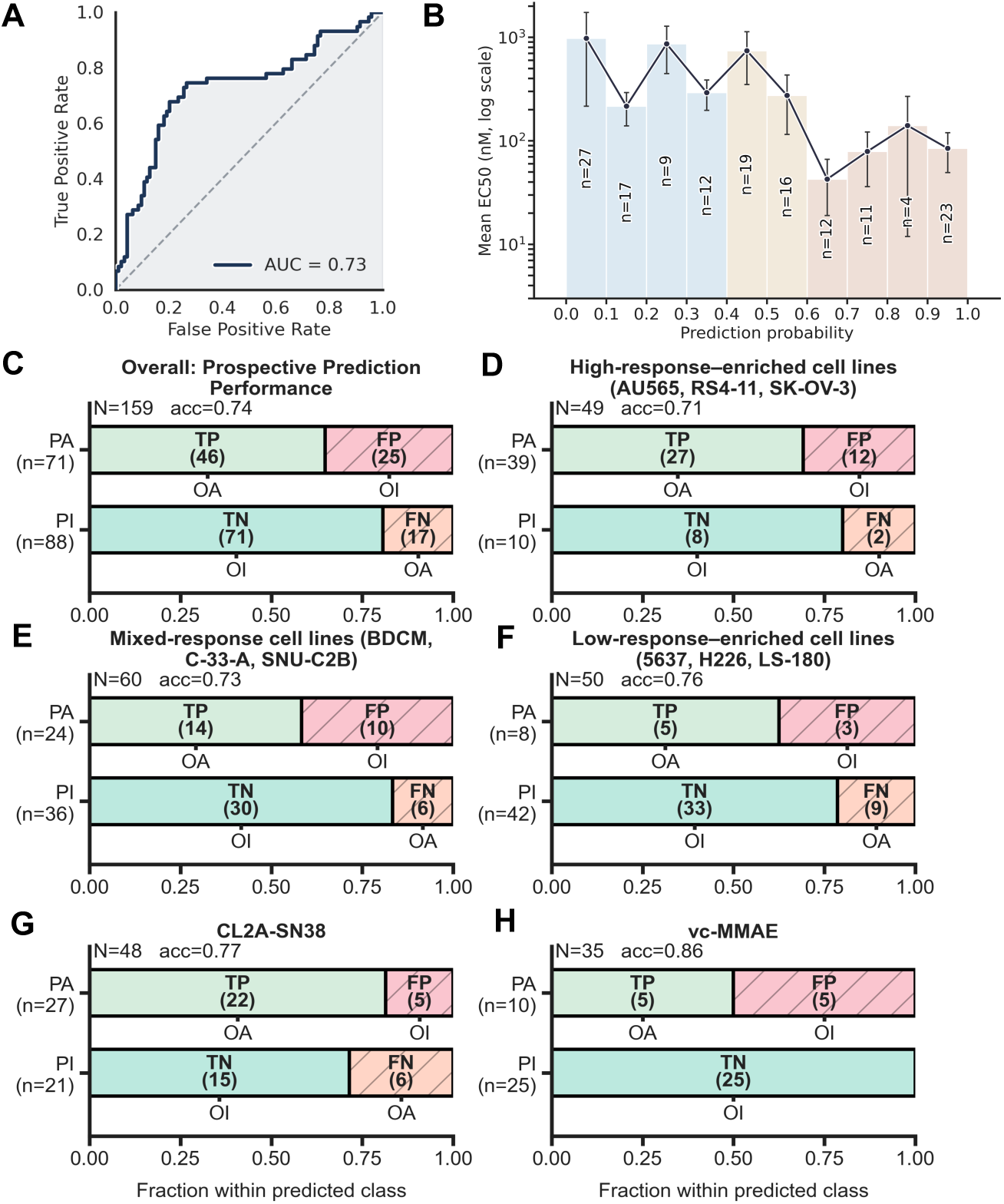
Prospective validation of AMM. (**A**) ROC-AUC performance for the AMM on 159-point prospective dataset. (**B**) Calibration analysis exhibiting the relationship between EC_50_ values and prediction probabilities, divided in low (light blue), mid (sand) and high (cream tan) tiers. (**C-H**) Confusion matrix analyses for overall and stratified performances, based on cell line tiers and CL2A-SN38 and vc-MMAE linker-payload systems.

Confusion matrix analysis at 0.5 probability threshold revealed sensitivity of 0.73 (95% CI; 0.61-0.82), specificity of 0.74 (95% CI; 0.64-0.82), PPV of 0.65 (95% CI; 0.53-0.75), and NPV of 0.81 (95% CI; 0.71-0.88), all reaching statistical significance (*p* < 0.0001; Fig. 5C). The LR of 2.81 indicates that a positive AMM prediction is nearly three times more likely to reflect true ADC activity than true inactivity. The high NPV is consequential for early ADC development, where a negative prediction carries 81% confidence that the combination is genuinely inactive, which can enable early elimination of failing designs and reducing the experimental burden on ADC programs.

To evaluate whether performance was consistent across the full AMM predicted probability range, the 159 combinations were stratified into cell line groups enriched with the likely sensitive (Fig. 5D), mixed (Fig. 5E), and resistant (Fig. 5F) predicted sensitivities. The sensitive-response-enriched tier (AU565, RS4-11, SKOV3) achieved sensitivity of 0.93 with two false negatives, meaning most active ADC design would be retained during an early screen. The NPV of 0.8 indicates that negative calls in this tier are highly informative (*p* = 0.0093; n = 49). The PPV of 0.69 indicates roughly two-thirds of positive predictions are correct, an acceptable trade-off in early discovery where missing a winning design could carry a greater cost than advancing a false positive for further testing. The mixed-response-enriched tier (C-33-A, BDCM, SNU-C2B), representing ambiguous predicted contexts and a rigorous biological test of the platform, achieved a sensitivity and specificity of 0.70 and 0.75, respectively, with a LR of 2.8 (*p* = 0.0017, n = 60). Critically, this demonstrates biological generalization rather than performance driven by extreme predictions at either end of the activity spectrum. The low-response-enriched tier (5637, LS-180, NCI H226) showed the platform’s strongest de-risking potential by specificity, at 0.92, with a LR of 4.3, where a positive prediction in these resistance-exhibiting model is over four-times more likely to reflect true activity (*p* < 0.0304; n = 50). This remains a useful confidence signal in the dataset for identifying unexpected ADC activity and in underrepresented cell lines that can be considered mimics of resistant disease.

A stratified analysis by linker-payload system revealed also significant performance for CL2A-SN38 (*p* = 0.0003; sensitivity 0.79, specificity 0.75, LR 3.1; n = 48; Fig. 5G) and vc-MMAE (*p* = 0.0008, sensitivity 1.0, specificity 0.83, LR 6.0; n = 35; Fig. 5H). The latter achieved zero false negatives, and a LR indicating that a positive prediction is six-times more likely to reflect true activity than inactivity. This was the strongest single performance signal in the prospective validation. GGFG-DXd also reached significance (*p* = 0.0080; sensitivity 1.0; PPV 0.37; NPV 1.0, LR 2.6; n = 38), whereas performance did not reach significance for GGFG-DXd (*p* = 0.0884; n = 38; Fig. S22A), with the high sensitivity and NPV indicating that AMM did not miss any truly active combinations. Performance did not reach significance also for vc-SN38 (*p* = 0.088; Fig. S22B), where a high PPV (0.80) alongside a low sensitivity (0.52), indicating that positive predictions were usually correct, but a substantial proportion of truly active combinations were missed.

Lastly, a stratified analysis by targeted antigen revealed that predictive performance generalized across multiple antigen-defined contexts rather than being driven by a single target class (Fig. S23). The strongest signal was observed for HER2 (*p* = 0.0002; sensitivity 0.76, specificity 0.86, LR 5.4; n = 38). Significant performance was also observed for CD22 (*p* = 0.0278; sensitivity 0.50, specificity 0.95, LR 10.0; n = 26), Nectin-4 (*p* = 0.0152; sensitivity 0.71, specificity 0.74, LR 2.7; n = 33), and Trop2 (*p* = 0.0164; sensitivity 0.58, specificity 0.86, LR 4.1; n = 33). CD22 showed a highly specific profile, whereas HER2, Nectin-4, and Trop2 showed more balanced discrimination across active and inactive ADC-cell line combinations. By contrast, performance did not reach significance for CD19 (*p* = 0.99; sensitivity 0.93; specificity 0.13; LR 1.1; n = 29; Fig. S23), where AMM classified 26 of 29 combinations as active. This high sensitivity coupled with low specificity indicates that AMM correctly identified CD19-targeted ADCs as broadly active in hematologic malignancy contexts, consistent with the strong CD19 biological signal evident in the prediction probability distributions (Fig. S14) but did not discriminate inactive CD19 combinations for solid tumor contexts.

### An open-source tool for predicting *in vitro* ADC efficacies

The ADC Design Platform is accessible through a secure online environment (www.adcpedia.com) that allows users to engage with both scientific resource and prediction elements. The “Open AMM-INVITRO Playground” portal is where users can select a target antigen, linker, payload, DAR, and generate predictions across the cell lines in ADCpedia. The interface allows linker-payload systems to be uploaded as encoded SMILES, SDF, SMI, or CSV formats or drawn-in using the JSME molecular editor.

## DISCUSSION

The central challenge in early ADC development is not a lack of chemical creativity – it is the inability to make reliable go/no go decisions before committing to the expensive and time-consuming process of testing across diverse tumor contexts. As a consequence, the field has optimized around what is feasible rather than what is optimal, defaulting to canonical linker-payload systems and well-characterized antigens, not because they represent the best biological matches, but because they reduce experimental uncertainty in the absence of better tools. The ADC Design Platform was built to address this selection problem directly, not by adding complexity to an already complex process, but by converting fragmented historical knowledge and incomplete biological data into actionable, prospectively validated predictions that scientists can use at an early and consequential decision stage in ADC development.

The platform’s most immediate practical output is its NPV of 0.81 (Fig. 5C). In the context of early ADC development, this means that when the platform predicts an ADC-cell line combination will be inactive, that call is correct 81% of the time. For an ADC program deciding which of dozens of designs to advance into costly *in vitro* and *in vivo* screening campaigns, the ability to confidently eliminate inactive combinations before a single experiment is performed represents a direct actionable reduction in experimental burden. This performance was validated across 159 ADC-cell line combinations spanning eight tumor types and five clinically validated target antigens, including three clinically approved linker-payload systems and, notably, most combinations which had no prior published ADC data. Collectively, this general performance demonstrates the platform’s potential to transition the current early ADC development process from empirical, to a rational more confident selection framework.

Antigen overexpression, both absolute and relative to normal tissues, govern ADC targeting of tumor cells. The ADC Design Platform suggested that cytotoxic outcomes are also determined downstream by processes beyond surface antigen density, including endocytosis, intracellular trafficking pathways, lysosomal processing, drug efflux capacity, oncogenic signaling and drug-responses – processes that are linked to the underlying biology of a tumor cell. The most direct evidence for this comes from the ablation studies and the platform’s significant predictive performance for CL2A-SN38-based ADCs, a linker-payload system that operates primarily through spontaneous hydrolysis rather than acidic conditions or through receptor-mediated internalization, functioning pharmacologically as a prodrug delivery system. Clinical data from the approved anti-Trop2 ADC sacituzumab govitecan, revealed no correlation between antigen expression and patient outcomes (*16*), and pharmacokinetic modeling demonstrates that circulating free antibody concentrations substantially exceed intact ADC levels throughout most of the dosing cycle (*15, 17*). Despite this non-classical mechanism, the AMM achieved statistically significant classification performance for CL2A-SN38 combinations, indicating that the platform learned key tumor cell biological features, particularly those that are relevant to topoisomerase I inhibition, including drug efflux and DNA repair capacity, which both affect SN38 therapeutic effectiveness (*51, 52*). Moreover, combining ABCG2 inhibitors with sacituzumab govitecan has been pursued (*53*). This finding also corroborates with two different ADC designs that target HER2 at the same epitope and with the same binding affinity produce markedly different results (*54*). The platform also achieved significant performance for the canonical protease-cleavable vc-MMAE- and GGFG-DXd-based ADCs, with vc-MMAE producing the strongest performance in the prospective dataset and GGFG-DXd showing high sensitivity that successfully captured all truly active combinations within that linker-payload class. Together with the CL2A-SN38 result, three of four prospectively evaluated linker-payload systems achieved statistically significant predictive performance spanning both classical receptor-mediated and prodrug-like cytotoxic mechanism. These findings generalize the platform’s utility beyond canonical internalization-dependent ADC designs and demonstrates that the architectural organization of AMM enabled it to capture cytotoxic determinants that are less mechanistically clear, nevertheless, effective.

Performance also generalized across antigen-defined contexts, with significant predictive performance achieved for HER2, Trop2, Nectin-4, and CD22 spanning both solid tumor and hematologic malignancy targets. The HER2 result represents the strongest antigen-specific signal, consistent with being the most extensively represented antigen in the training data. The CD22 result is particularly informative achieving a positive LR ratio of 10.0 with high specificity, indicating that the platform can confidently identify inactive CD22-targeted ADC combinations not only for hematologic malignancies but also for solid tumors. The poor performance for CD19 indicate the model would benefit from expanded training data for CD19 for additional tumor types.

This is not definitive proof as there are important pharmacokinetic and biodistribution parameters in patients that are critical for determining ADC efficacy (*55, 56*). The work presented here stops at *in vitro* and is not designed to perform past early preclinical ADC development. Nevertheless, it provides a biologically grounded comprehensive ML-based system to support these known clinical observations.

The prospective validation design was constructed to circumvent the common limitation of ML models in drug discovery that performance metrics reflect pattern recognition within known data rather than generalizability. Most of tested cell lines had no prior published ADC data, meaning the platform was operating in previously unchartered biological territory for the majority of its predictions. We further evaluated combinations with widespread probabilities across all three design dimensions (targeted cell line, linker-payload system, targeted antigen) rather only combinations likely to yield favorable outcomes. This was a key stress-test as it challenged the platform against a wide spectrum of biological scenarios an ADC program would most likely encounter.

We could not find any publicly accessible prospective platforms to compare against. The industry benchmark against which the AMM’s prospective performance was evaluated was derived from Sturm et al. (*50*). This study conducted a large-scale assessment of deep learning models trained on public small molecule bioactivity data and transferred to internal pharmaceutical industry datasets from AstraZeneca and Janssen. In that study, models were built on the ExCAPE-ML dataset covering 526 unnamed heterogeneous protein targets and nearly 50 million quantitative structure-activity relationship datapoints, which uses structural fingerprint descriptors alone as embedded features. The prospective evaluation consisted of applying those models to novel compounds tested against the same protein targets represented in the training data. Under these conditions, the prospective performance dropped 13-21% compared to the retrospective cross-validation performance, which was similar to what was observed in our retrospective evaluation (Fig. 3E). Although, these prospective and retrospective results are similar, our study has three noticeable differences. First, the drug-target context is more complex as ADCs are multicomponent and biological-small molecule hybrid drugs whose cytotoxic outcome depends on the interplay between these components and the underlying cellular components. Second, AMM was trained on 1,167 curated datapoints, orders of magnitude fewer than the millions available in ExCAPE-ML, suggesting the large-scale pretrained biological embeddings were instrumental in driving performances. Third, unlike inhibition assays, AMM’s prospective validation is a whole-cell cytotoxic outcome integrating biological processes from antigen binding through apoptosis. Moreover, the majority of predictions were on cell lines with no previous ADC history. Along with additional multi-omics information, this suggests that AMM can profile the unseen cell lines into a biologically meaningful representation to predict for ADC activity where there was previously no record for these cell lines. Taken together, the ADC Design Platform predicted ADC activity at industry standards but in a biologically rich and more complex environment.

The platform’s utility for broad-problem applications in ADC development was further demonstrated by the cell line selection workflow. This was an unbiased method of ranking cell lines in ADCpedia by predicted sensitivity probability across antigen and linker-payload combinations. This provides ADC scientists with a principled, data-driven approach to prioritizing which tumor contexts to test first and target or avoid and has the potential to significantly aide the current practice of defaulting to the best-characterized cell lines regardless of their biological relevance. This is especially important for emerging and less-characterized antigens where preclinical and clinical literature is limited or non-existent. Indeed, this work exhibited GENCEP-imputed protein expression profiles for relevant target antigens across 98 cancer types, providing a standardized biological atlas that did not previously exist. Additionally, this ability extends to any protein of interest and multiple proteins can be coupled to obtain a comprehensive atlas interconnected with past ADC designs and cytotoxic outcomes. This provides an effective resource that can be used to hone precision when utilizing AMM for predictive work.

Several limitations of the current platform should be acknowledged. GENCEP’s protein intensity predictions were validated against ground truth values with R^2^ = 0.75 and 0.86 for ADC target antigens and all proteins, respectively (Fig. 2F and S7). These are strong but imperfect correlations. Biochemical validation assays could provide the necessary validation through methods such as flow cytometry or Western blot. However, flow cytometry would only capture cell surface expression, whereas GENCEP models whole-cell protein abundance from lysate-based proteomics, and Western blot quantification lacks the precision of mass spectrometry-based intensities. The current platform is optimized for binary classification at the clinically relevant 10 nM EC_50_ threshold. For programs pursuing highly potent sub-nanomolar ADCs, AMM may have reduced precision in distinguishing activity gradations within that activity range. The platform’s binary classification framework also does not distinguish between canonical receptor-mediated cytotoxicity and prodrug-like payload release mechanisms, including any non-receptor-mediated opsonization. However, this may be a feature rather than a limitation, as the AMM was trained on composite cytotoxic outcomes and its significant performance for CL2A-SN38 demonstrates it can predict tumor cell sensitivity to SN38-specific release through the biological integration employed in this work. However, future iterations incorporating mechanism-aware features may further improve specificity for individual linker-payload classes and for specific relapse/refractory disease states. As described, antibody amino acid sequence was scarce. In the future, as more antibody-specific information is collected, incorporating binding specificity will be important as this can greatly affect internalization pathways and kinetics (*5*). Most critically, the platform does not yet account for *in vivo* pharmacokinetics, tumor uptake, bystander killing effects, or immunological contributions to ADC efficacy, all of which strongly influence the translation from preclinical to clinical outcomes. Its value is therefore as an early-stage decision support tool that narrows the experimental search space before translational commitments are made and, not as a substitute for the full preclinical development pipeline. Finally, the platform faces the broader challenge in ML-based drug discovery programs, which is insufficient high-quality standardized data (*57*). Most ADC records had no or minimal associated activity information (Fig. 2A), design components were narrow (Fig. 2D), and cytotoxicity conditions varied substantially across sources (Fig. 4B). In-house standardized data generation remains the gold standard and, therefore, the cytotoxicity assay approach used in this study can be a guide.

The ADC Design Platform was not devised to substitute how scientists develop new ADC drugs. Instead, it enables them to advance more effectively by providing increase precision for early selection in a much more comprehensive manner. Instead of beginning with a canonical cell line, an estimated antigen surface density, and a familiar linker-payload combination, the platform enables first decisions to be made against a quantified probability of activity across the biological landscape most relevant to a given ADC program – whether that program targets a well-characterized antigen through canonical delivery mechanism or an emerging antigen through a non-canonical delivery mechanism. The historical ADC development record reflects the cumulative cost of incomplete information at early evaluations, and the platform’s overall performance demonstrates that a different approach is possible, one where the question “which ADC design is most likely active in which tumor context?” can be answered with quantified, biologically rooted confidence before a single experiment begins. That is not a promise of clinical success, but it is a potential tool for getting ADC programs past a critical early resource-intensive bottleneck faster, with better biological justification, and with greater confidence in the decisions that follow.

## Material and Methods

### Database building

Although previous curations of ADC structure-activity databases were created by extracting as much information as possible, these inherently leave large gaps. Databases built on searching multiple patent organizations and extracting information from the R&D pipelines listed on the websites of pharmaceutical companies (*2*) leads to significant inferring of structure and/or activity information for a given ADC or leaves gaps in the database, a concern that is detrimental to training (*57*). Additionally, such extensive searches most likely identify ADCs developed very early in the field and are not realistic into today’s much more refined field.

The curation process for ADCpedia focused on preclinically and clinically validated ADCs listed in the ADC DrugMap (*43*). Surprisingly, ADC DrugMap structures included several molecules that do not traditionally constitute an ADC such as biparatopic or bispecific antibodies and ɑ-particle emitting payloads (*43*). To avoid a potential ‘data swamp,’ ADCpedia was refined based on standardized inclusion criteria, that is, considering only full monospecific IgG antibodies, hetero-bifunctional chemical linkers, and traditional small molecule/anti-mitotic peptide payloads, akin to chemotherapeutic activities. A targeted search was then performed on these standardized designs in the literature between January 1, 2000, to April 15, 2024.

A critical second inclusive criterion was that structural information must be accompanied by cytotoxicity EC_50_ values, based on consistent methodologies for determining quantitative EC_50_ values. Therefore, only studies reporting cytotoxicity assays conducted in multi-well plate formats, single time-point incubation periods (*e.g*., 72 h, 120 h, etc.), ADC concentration-response relationships, and readouts based on fluorescence- or luminescence-based reagents were included in the database. As a result of these standardization measures, a total of 151 articles and patents were collected and procured a total of 1,167 datapoints on ADC linker-payload compositions, antibody conjugation sites, DARs, the accompanying EC_50_ values, target antigen, tumor types, and tumor cell line names.

### ADC and tumor cell line embeddings

Physicochemical features for the payload and linker structures were computed using the RDKit cheminformatics (*58*) and Descriptastorous libraries (*59*). A set of 200 physicochemical properties was calculated, including log P, molecular weight, hydrogen bond donors and acceptors, rotatable bonds, and topological polar surface areas. Additionally, MACCS fingerprints (167 bits) were also computed (*58*), thus also encoding the antibody conjugation site (lysine, cysteine, glutamine) via terminal linker chemistry. DAR was added as a distinct feature. Payloads were stratified into the three mechanisms driving cytotoxicity, namely, DNA damagers, microtubule disruptors, and topoisomerase I inhibitors, and one-hot encoding was used to assign the correct mechanism to each payload.

RNA-seq profiles ‘read counts’ for approximately 37,600 protein encoding genes from 1,479 human tumor cell lines that represented 98 distinct tumor types were obtained from the Cell Model Passport (*44*) (Fig. S1), including 12 non-cancer cell lines representing diverse tissue types. For ADC-relevant tumor cell lines, features were computed using two approaches. First, the canonical amino acid sequences for target antigens (no alternatively spliced isoforms) were retrieved from UniProt and transformed into 1,280 Evolutionary Scale Modeling (ESM; 650M model) embeddings (*60*). Principal component analysis (PCA)- was then applied, preserving 95% of the variance and resulting in 356 dimensions. The antibody amino acid sequences were intentionally not considered and rather treated as a constant feature because to the best of our search, limited publicly available sequence information was found, most likely for proprietary reasons. Second, to address the aberrant activity of relevant intracellular processes related to ADC performance (*5, 7, 61*), the transcriptomic data from the Cell Model Passport repository (*44*) was utilized to derive transcriptomic signatures for all cell lines in ADCpedia. Specifically, the entire raw read counts of mRNA expression profiles for the human tumor cell lines, totaling 37,603 protein-encoding genes, were integrated into ADCpedia. The cell lines with published ADC activities were then subjected to PCA formatting to reduce dimensionality while retaining 95% of the variance, resulting in 910-dimensional cell line embeddings geared for ML applications. Cancer model names from the Cell Model Passport were used to augment the database with detailed annotations on tissue, cancer type, and sample site for each included cell line.

### GENCEP model

The GENCEP model took as inputs (i) raw mRNA expression data, (ii) ESM embeddings of all protein sequences, both PCA-reduced, and (iii) gene-specific read counts for cell lines (Fig. S2B). ESM embeddings are processed through a fully connected neural network with two layers, each containing 512 neurons. The mRNA data was processed through two dense layers of 512 neurons each. The read counts were passed through an additional dense layer with 64 neurons The three outputs were then concatenated into a 1,088 vector and fed into a network consisting of three fully connected layers with decreasing neuron counts (1,024, 512, 64). ReLU activation function, batch normalization, and a dropout of 15% were applied to every dense layer. The outputs from the mRNA expression and read count components were also utilized as embedded inputs for the AMM model (described in the next section). The model was trained using Root Mean Squared Error (RMSE) loss to measure the difference between predicted and experimental protein intensities:

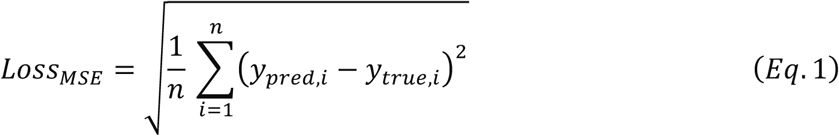

Optimization was performed using the Adam optimizer (*62*) with an initial learning rate of 1E−3 and a weight decay of 1E−5. Dataset split was performed with the goal of evaluating the applicability of the model to ADC-target antigens. Thus, all the antigens in ADCpedia were kept in the hold-out test set, which was further populated via random sampling to a size equal to 10% of the dataset. Training and validation sets were also built via random sampling (80% and 10%, respectively). Early stopping was implemented based on the validation RMSE value with a patience of 10 epochs.

### AMM model

#### Data preprocessing

A StandardScaler (*63*) was applied for continuous features. For labeling, EC_50_ values were binarized at 10 (EC_50_ < 10 nM assigned label 1; otherwise, 0). To prevent data leakage, we performed Butina clustering (*64*) (80% threshold) on the Tanimoto dissimilarity matrix of the linker-payload fingerprints. The resulting clusters were then partitioned with stratification into training, validation, and testing sets in proportions of 80%, 10%, and 10%, respectively. Additionally, the training was balanced by oversampling the minority class (0).

#### Architecture

The architecture was a multimodal CNN that processed the following inputs through three streams, Stream 1: (i) DAR and intracellular target information, (ii) PCA-reduced ESM embeddings of the target antigen Stream 2: (iii) Cell line-specific mRNA embeddings (from the GENCEP model), (iv) mRNA read count embeddings for the target antigen from GENCEP and normalized mRNA raw counts for mode-of-action (MOA)-relevant proteins: drug-efflux transporters (ABCB1, ABCG2), endosomal and lysosomal trafficking/activity genes (RAB7A, CTSB, LAMP1), and microtubule-associated payload response genes (TUBA1A, TUBA1C) (*65–69*); DNA-damage response and genomic stress genes (BRCA1, BRCA2, TP53, RAD51, CHEK1, CHEK2, CDKN1A), PI3K/AKT/mTOR pathway genes (PIK3CA, AKT1, MTOR), anti-apoptotic cell-survival genes (BCL2, BCL2L1, MCL1), and hypoxia-associated genes (HIF1A, CA9) (*70–77*); and detoxification and oxidative-stress genes (GSTM1, GSTP1, NQO1) and epigenetic regulators linked to tumor plasticity and treatment response (EZH2, KDM5C) (*78–84*), (v) predicted antigen expression intensities for antigen and MOA-relevant proteins (from the GENCEP model). Stream 3: (vi) physicochemical (200 total) and MACCS fingerprint descriptors (167 bits) of the payload-linker system (Figure S1A).

Each feature stream in the multimodal framework first underwent an initial linear projection to reduce dimensionality (Figure S2A). Each raw feature block was first passed through a feature-specific linear projection layer, which either reduced, preserved, or expanded its dimensionality. The linearly projected features within each stream were concatenated and passed to a dedicated 1D convolutional block, whose layers, hidden channels, kernel size, and dropout were tuned via Optuna. All convolutional outputs then went through a global average pooling step and were subsequently concatenated into a combined embedding, which was refined by an attention mechanism. Finally, a fully connected classifier (with ReLU activations, batch normalization, and dropout) output a sigmoid probability for binary classification.

A custom binary cross-entropy (BCE) loss function was designed to emphasize classifications far from the selected nM threshold boundary while down-weighting ambiguous samples near the 10 nM threshold. Specifically:

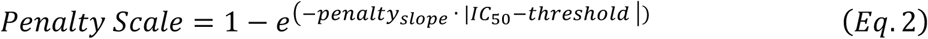

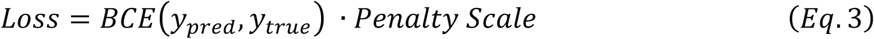

where *BCE*(*y*_*pred*_, *y*_*true*_) was the standard binary cross-entropy loss, *penalty_slope_* controlled the rate at which the penalty decreased near the boundary (set to 10), EC_50_is the quantitative value for each data point, and *threshold* was set to 10 nM. Early stopping was implemented based on validation loss, halting training if no improvement was observed over eight consecutive epochs. Learning rate scheduling via ReduceLROnPlateau (*85*) was employed to reduce the learning rate when validation loss plateaued.

#### Hyperparameter optimization

Hyperparameter searches were conducted using the default Tree-structured Parzan Estimator within Optuna (*86*). The exploration covered 1 to 3 CNN layers, hidden dimensions ranging 32–512, kernel sizes up to 7, dropout up to 50%, learning rates in [E-5, E-2], and batch sizes of 32, 64, or 128, each with an early-stopping patience of 10. The 10 nM decision threshold was used for labeling. The best identified hyperparameter combinations are detailed in Table S1.

#### Ablation studies

To quantify the contribution of each input modality, ablation was performed on the AMM model’s features. In each ablation, feature branches were removed from both the data pipeline and the model graph, such that the ablated model was retrained using only the remaining inputs. The feature branches encompassed target antigen ESM embeddings, predicted antigen expression intensity, antigen mRNA count embeddings, predicted MOA-relevant protein intensities, MOA-relevant gene mRNA counts, learned cell line embeddings, chemistry descriptors (RDKit features and MACCS fingerprints), as well as DAR and intracellular target. Each feature-ablated model was selected based on the best validation ROC-AUC. Performance loss was summarized as the difference between validation accuracy between the full model and the ablated one.

### Computational infrastructure

All computational tasks were performed on high-performance computing clusters provided by the Digital Research Alliance of Canada, including a Béluga virtual machine allocation with 8 virtual CPUs and 30 GB of RAM (p8-30gb configuration). The operating system used was Ubuntu-22.04.4-Jammy-x64-2024-06. For inference and website integration, job scheduling and resource allocation were managed by the SLURM workload manager. The computational environment was also configured to support multi-threaded operations and scale with demand.

### External blinded test set

The external ADCs were kindly provided by Dr. Mark Barok (University of Helsinki). ADC structures (antibody name, linker type, payload, and DAR) and the cell lines tested against were provided. Activity classes were determined by AMM and sent back to Dr. Barok for verification. A portion of the ADCs has since been published (*87*). Chemical structures and activity data for this set are reported in Table S3. The cytotoxicity assay procedures were not provided.

### Prospective validation workflow

#### Cell lines and cell culture procedures

Cell lines were procured from Cedarlane Labs (Burlington, Ontario, Canada), the Canadian distributor for American Type Culture Collection (ATCC). Frozen tubes were thawed and established according to ATCC-recommended protocols for each individual cell line. Cultures were expanded until sufficient cell numbers permitted cryopreservation of working stocks. Original thawed cell cultures were used for cytotoxicity testing across no more than 5-10 passages, corresponding to approximately 1-2 months of continuous culture. Cell viability was assessed routinely by Trypan Blue exclusion using manual cell counting and a TC20 Automated cell counter (Bio-Rad). Cytotoxicity assays were only accepted when cell viability was ≥85%.

#### Antibodies, linker-payloads, and ADCs

All antibodies and linker-payload systems were purchased from MedChemExpress using Cedarlane Labs as the Canadian distributor. The catalog numbers the antibodies were: trastuzumab (HY-P9907), sacituzumab (HY-P99045), enfortumab (HY-P99016), loncastuximab (HY-P99711), and inotuzumab (HY-P99264). The catalog numbers for linker-payloads were: CL2A-SN38 (HY-128946), vc-SN38 (HY-131057), GGFG-DXd (HY-13631E), and vc-MMAE (HY-15575). Trastuzumab deruxtecan (HY-138298A), free SN38 (HY-13704), rituximab (HY-P9913), and human IgG1 kappa isotype (HY-P99001) were also purchased for cytotoxicity assay optimization and controls and drug release experiments.

#### ADC construction

Bioconjugations were performed according to protocols established in seminal studies for each linker-payload system (*14, 88, 89*). Briefly, each antibody (2 mg) was reduced by incubation with 74-fold molar excess of dithiothreitol (DTT) in a 5.4 mM EDTA, 1x PBS (pH 7.4) for 70 minutes at 37 °C, with gentle agitation every 15 minutes to maximize interchain disulfide reduction. Reduced antibody was buffer-exchanged into 75 mM sodium acetate, 1 mM EDTA (pH 6.5) using Amicon Ultra centrifugal filters (100 kDa MWCO; 500 µL capacity, MilliporeSigma) by sequential centrifugation at 10,000 x g for 5 minutes, repeated seven times to a final retentate volume between 50-100 µL. Linker-payload stocks prepared in DMSO were added to the reduced antibody at molar excesses ranging from 2- to 30-fold, depending on the target DAR, with the addition volume not exceeding 15 µL per 0.5 mL of reduced antibody solution to minimize solvent perturbation. For GGFG-DXd constructs, the linker-payload molar excess was capped at 15-fold to avoid over conjugation or antibody denaturation. Conjugation reactions proceeded for 40 minutes at room temperature under constant rotation and protected from light. The resulting ADC was buffer exchanged back into 1x PBS (pH 7.4) as described above. DAR was determined immediately by UV-Vis spectrophotometry and protein concentration quantified (NanoDrop 2000; ThermoFisher). Each ADC was aliquoted at 50 µg for single-use and stored frozen at -20 ℃ until cytotoxicity testing.

#### DAR quantification and ADC uniformity

DAR was determined for all ADC constructs immediately following conjugation by UV-Vis spectrophotometry and represented the primary DAR characterization at time of cytotoxicity testing. Extinction coefficients for each antibody were determined experimentally by serial dilution in PBS at the relevant measurement wavelengths. Extinction coefficients for each linker-payload were obtained from the published literature. The DAR for each construct was calculated using the following formula:

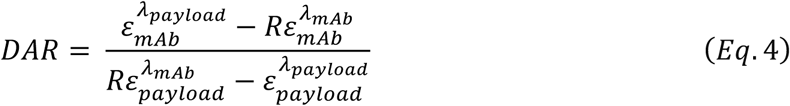

Where:

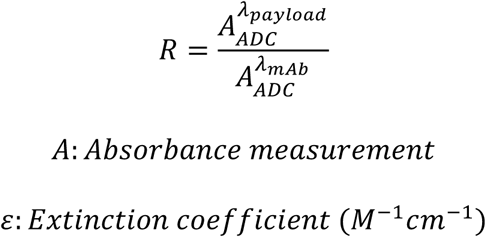

The corresponding wavelength (λ) for each extinction coefficient and absorbance measurement, specified via superscript. Subscripts specify the molecular species being referenced (e.g., antibody, payload, or ADC) for each absorbance or extinction coefficient term. Extinction coefficients used for all constructs is provided below In Table 1.

**Table 1.** Extinction coefficients.

| **Antibodies** | **$\epsilon_{280}$** | **$\epsilon_{269}$** | **$\epsilon_{259}$** |
| --- | --- | --- | --- |
| Inotuzumab | 232340 | 188828 | 115581 |
| Enfortumab | 246393 | 200472 | 121310 |
| Trastuzumab | 227920 | 215044 | 134976 |
| Sacituzumab | 218162 | 176897 | 109715 |
| Loncastuximab | 246258 | 199658 | 119823 |
| **Linker-payloads** | **$\epsilon_{280}$ (drug)**<br>**(literature values)** | **$\epsilon_{248}$ (drug)** | **$\epsilon_{380}$ (drug)** |
| vc-MMAE | 1500 | 19992 | - |
| GGFG-Dxd | 5000 | - | 20589.9 |
| CL2A-SN38 | 6100 | - | 20772.4 |
| vc-SN38 | 6100 | - | 14935.8 |

#### ADC integrity and homogeneity

Size-exclusion chromatography-high performance liquid chromatography (SEC-HPLC) was performed on nine representative ADC constructs drawn from available frozen stocks using a TSK Gel G3000SWXL column (Tosoh Biosciences, Cat. No. 08541; 7.8 mm x 40 cm, 5 µm particle size) on an UltiMate 3000 Dionex HPLC system, monitored at 280 nm. An isocratic mobile phase comprising 5% isopropanol in 0.4 M NaCl, 0.015 mM phosphate buffer (pH 7.4) was used at a flow rate of 0.9 mL/min. Samples were diluted to approximately 100 µg/mL and 15 µL was injected. The majority of constructed ADCs were monodispersed with minimal aggregation across all four linker-payload systems (Fig. S15A-D). Two prepared ADCs showed 13% and 15% higher molecular weight species and most likely due to long-term storage. Two UV-Vis DAR values were revised following SEC analysis but remained within the DAR range used for prospective AMM predictions and did not alter the classification of any ADC-cell line combination. Native SEC-mass spectrometry (nSEC-MS) was used to evaluate an additional eight representative ADC constructs (Fig. S15E-H). Prior to analysis, samples were deglycosylated overnight with PNGase F (Sigma, Cat. No. F8435-300UN) at 37 ℃. Antibodies/ADCs were subsequently diluted to 1 mg/mL, injected and separated with an ACQUITY UPLC Protein BEH-SEC column (200 Å, 1.7 µm, 2.1 mm x 150 mm) and an isocratic mobile phase (50 mM Ammonium acetate) at 100 µL/min using an UltiMate 3000 Dionex HPLC system. Samples were electrosprayed into an Orbitrap Exploris 480 operated in positive ion mode (3400V) scanning from 2500-8000 m/z in MS mode with normalized AGC of 300% and a collision energy of 120. Chromatograms were anlayzed with Thermo Xcalibur Qual Browser (v4.4.16.14). ADCs were monodispersed (∼5-6 min) with no evidence of aggregation. Small molecule contaminants eluted at 7-8 min.

### *In vitro* cytotoxicity assay standardization

#### Cell density and incubation period

Cytotoxicity assay parameters were benchmarked against the diversity of protocols represented in ADCpedia prior to any prospective experimental testing. Seeding density and incubation duration were extracted for each record and plotted against corresponding EC_50_ potency category: active (<10 nM; deep purple), intermediate activity (10-100 nM; purple), and inactive (>100 nM; light purple), which record frequency represented by circle size (Fig. 4B). Incubation periods ranged from 24 to 120 hours. From this analysis, a seeding density of 5,000 cells per well was selected as a standardized threshold, chosen to avoid any potentially artificially inflated potency readings with lower cell densities. A 72-h incubation period was established as the standard exposure time, reflecting the modal value across ADCpedia records (Fig. 4B). Both parameters were fixed uniformly across all nine cell lines and all four linker-payload systems for the entirety of the prospective validation.

#### ADC concentration range and EC_50_ calculation

The ADC concentration range was determined from a systematic survey of ADC cytotoxicity records in ADCpedia, evaluating the fold-range and number of concentration points reported across the literature, a few referenced here (*12, 14, 21, 90*). The most common concentration-fold range across records was approximately 62,000 – 100,000-fold, with 8- or 9-point concentration series. Based on this, a fixed 8-point concentration series was established: 0.008, 0.04, 0.2, 1, 5, 26, 130, and 650 nM, representing an 81,250-fold range.

EC_50_ values were calculated by fitting concentration-response data to a four-parametric log-logistic model (4pLL-model) in GraphPad Prism (Version 11.0.0), with lower and upper asymptotes constrained to 0 and 100, respectively, consistent with the assumption that solvent control wells represent 100% cell viability and maximal drug effect converges to 0% viability. This fitting approach follows the optimally designed protocol for *in vitro* cytotoxicity experiments described by Schürmeyer *et al.,* (*91*). Concordance between Prism-derived EC_50_ values and those obtained by independent software implementations of the 4pLL model was confirmed by comparison against EC_50_ values from Schürmeyer *et al*. and Proctor *et al*. (*91, 92*), for well characterized small molecule cytotoxic agents tested against the same cell lines (Fig. S16). This validates the Prism 4PLL method for implementation for all subsequent ADC cytotoxicity calculations.

The concentration range and EC_50_ calculation method were validated using the approved ADCs T-DM1 and T-DXd incubated with SKOV3.ip cells (kind gift from B. Chabot, Université de Sherbrooke; Fig. S13). SKOV3.ip is an aggressive and invasive subline derived from the parental SKOV3 ovarian cancer cell line, expressing approximately 2-fold higher HER2 surface density than the parental line (*93*). As SKOV3.ip is not represented in ADCpedia, it served as a genuinely independent validation cell line for assay optimization without risk of introducing selection bias into the prospective validation. Reproducible EC_50_ values were obtained on both sides of the 10 nM activity threshold across triplicate technical and biological replicates, confirming that our selected concentration range captured the full dynamic range of ADC cytotoxic activity relevant to AMM binary classification. The standardized conditions of 5,000 cells per well, 72 h incubation, and 8-point concentration range were subsequently applied uniformly across all prospective ADC-cell line combinations without further cell line-specific tuning, preserving a real-world experimental context in which the platform’s generalizability could be attributed to AMM predictions as opposed to condition-specific optimizations.

#### ADC specificity evaluation

To characterize the cytotoxic mechanism of representative ADC constructs, specificity was evaluated using HER2 positive SKOV3.ip and HER2-negative CHO-K1 cells (Cedarlane Labs) as a matched antigen-positive and antigen-negative pair. First, the naked antibodies used for the ADCs in the prospective evaluation demonstrated no measurable cytotoxicity on AU565 or RS4-11 (sensitive) cells, with EC50 values >1000 nM or not calculable. This confirmed that the antibody component contributed no cytotoxic activity independent of payload delivery (Fig. S18).

#### SN38 release kinetics

Concentrations of SN38 release from T-CL2A-SN38 and T-vc-SN38 at physiological and acidic pH were measured by ultra-performance liquid chromatograph-quadrupole time-of-flight high resolution accurate mass spectrometry (UPLC-QTOF HRMS) using a Xevo G2-XS instrument and UPLC I-Class system (Waters, Milford, MA) at the Université de Montreal, Faculté de pharmacie, Plateforme de biopharmacie, *Solutions en AME-Tox*. Instrument parameters including cone voltage and collision energy were optimized by direct infusion of SN38. SN22 was used as an internal standard at a working concentration of 2,000 ng/mL in methanol-water (70/30) containing 0.1% formic acid.

A stock solution of SN38 was prepared at 5 mg/mL in DMSO and serially diluted in methanol-water (30:70) containing 0.4% glacial acetic acid to generate calibration standards at 2.5, 5, 10, 25, 50, 100, 250, 500, 1000 and 2000 ng/ml. Samples (10 µL) were diluted 20-fold in the same diluting solvent (190 µL) and 10 µL of internal standard working solution was added to 200 µL of each calibrator and sample prior to gentle vortex mixing. Chromatographic separation was performed on a Waters Acquity BEH C18 column (50 x 2.1 mm, 1.8 µm) using gradient elution at 0.4 mL/min over a total acquisition time was 4.0 min. Data were acquired in TOF-MSe mode. Calibration curves were constructed using the analyte-to-internal standard peak area ratio with quadratic regression and 1/x weighting. Free SN38 concentrations were determined in ng/mL and converted to µM by dividing by the molecular weight of SN38 (392.4 g/mol). Molar percent SN38 released was calculated by dividing the experimentally determined free SN38 concentration by the theoretical SN38 concentration based on the construct DAR and total antibody concentration.

To characterize the spontaneous payload release properties of the CL2A-SN38 and vc-SN38 linker-payload systems under conditions relevant to in vitro cytotoxicity assays, UPLC-QTOF HRMS was used to measure SN38 release at physiological pH 7.4 and acidic pH 4.0 over 72 h for both free linker-payload and trastuzumab-conjugated ADC constructs.

### Online interface

Allocation and deployment with Digital Research Alliance of Canada involved using a virtual machine allocation for public access, facilitated by a “Floating IP.” A subdomain (server.adcpedia.com) was linked to this IP, and the application was developed with WordPress for the frontend and Django for the backend, integrating prediction models and data pipelines. Nginx and Gunicorn were used for deployment, while SSH managed the server. Project files were transferred, and settings were configured to handle cookies, CSRF, and CORS. Security rules for HTTP and HTTPS enabled requests on ports 80 and 443. Nginx was set up to redirect traffic to Gunicorn, serving the Django application. Static files were collected, database migrations executed, and SSL certificates obtained via Certbot for HTTPS security. File permissions ensured proper access, and multiple Gunicorn instances were able to maintain performance under heavy loads. Monitoring of Nginx logs helped detect and resolve issues. CSRF tokens secured POST requests, while CORS was configured to manage requests from different origins, essential for the Digital Research Alliance of Canada deployment.

### Statistical analysis

Multivariable analytical methods were used to generate associations between ADC components and antigen expressions and EC_50_ values. Continuous variables were compared using Pearson correlation coefficients and linear regression models, with model performance quantified by the coefficient of determination (R^2^). For instance, correlations between scaled mRNA read counts and true protein intensities were assessed by scatter plots overlaid with regression lines, while differences in antigen expression across predefined EC_50_ bins were examined using boxplots and strip plots. Outliers were identified and excluded based on Z-score and interquartile range criteria with the box encompassing the 75^th^ interquartile range (IQR) and the mean indicated by horizontal lines in the boxes. Box whiskers span the 25^th^ IQR. The IC₅₀ values were log-transformed (pEC₅₀ = −log₁₀[EC₅₀]) when appropriate. In addition, classification models for drug sensitivity (using an EC₅₀ threshold of 10 nM) were evaluated by ROC-AUC analyses for validation, test, blind, prospective datasets. Confusion matrices were further constructed for overall performance for generalizability, across cell group tiers, linker-payload systems, and certain cell line–specific analyses, generate statistical significance defined as *p*<0.05, sensitivity, specificity, positive predictive value, negative, predictive value, and likelihood ratio. Data was analyzed using Graphpad Prism and/or Python (pandas, SciPy, scikit-learn, and seaborn) for figure presentation. Concept figures were generated using BioRender.

## Acknowledgments

This research was funded by the Canadian Institutes of Health Research, the Natural Sciences and Engineering Research Council of Canada, and internal funding by the Faculty of Medicine, University of Ottawa. The authors thank the Digital Research Alliance of Canada for computational resources.

## Author contributions

https://credit.niso.org/ Conceptualization: HM, FG, JVL

Methodology: HM, RT, JGS, MGW, MN, GC, HS, ME, TY, OAB

Funding acquisition: FG, JVL

Project administration: FG, JVL

Supervision: FG, JVL

Writing – original draft: JVL

Writing – review & editing: HM, FG, JVL

## Data and materials availability

All data is included in the supplementary materials. The codes, database, and model architecture are available at https://github.com/giaguaro/ADCpedia. Unlimited access for performing predictions is available at the following link: https://molcomp.com/models/adcpedia/adcpedia-predictions.

