## Supplementary Material for "Interconnecting ADC Structure with Tumor Cell Biology with Multimodal Learning"

**The PDF file includes:**

Supplementary Text
Figs. S1 to S23
Tables S1 to S6
Supplementary References

### SUPPLEMENTARY TEXT AND FIGURES

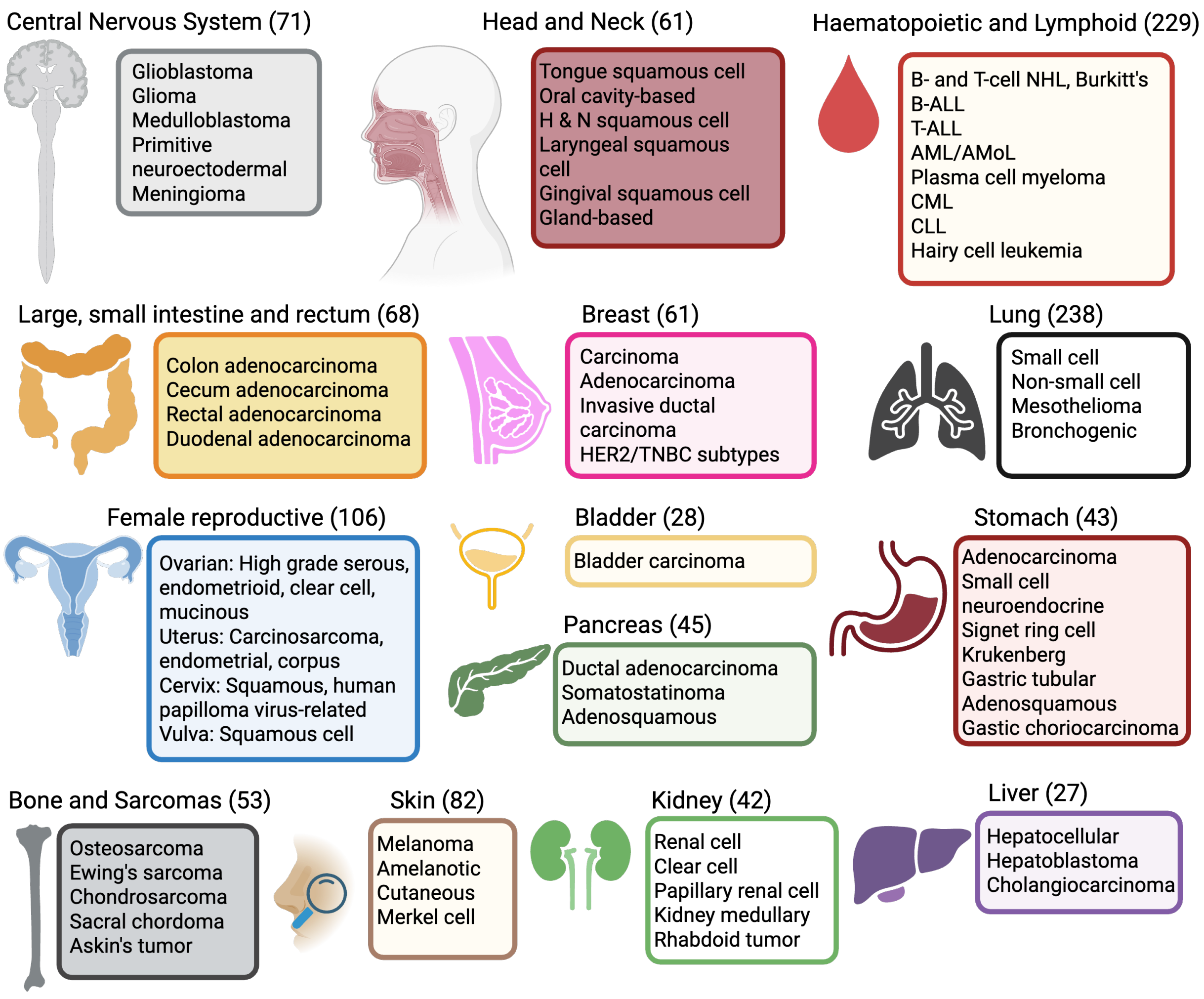

***Fig. S1. Tumor cell lines and tumor type as part of the multimodal framework of the ADC Design Platform.***

*An overview of the cell lines representing diverse organ or physiological systems, cancers, and specific subtypes. The approximate number of cell lines for each system at the time of platform building is in parentheses. Specific cell lines and their omics profiles are detailed at cellmodelpassports.sanger.ac.uk.*

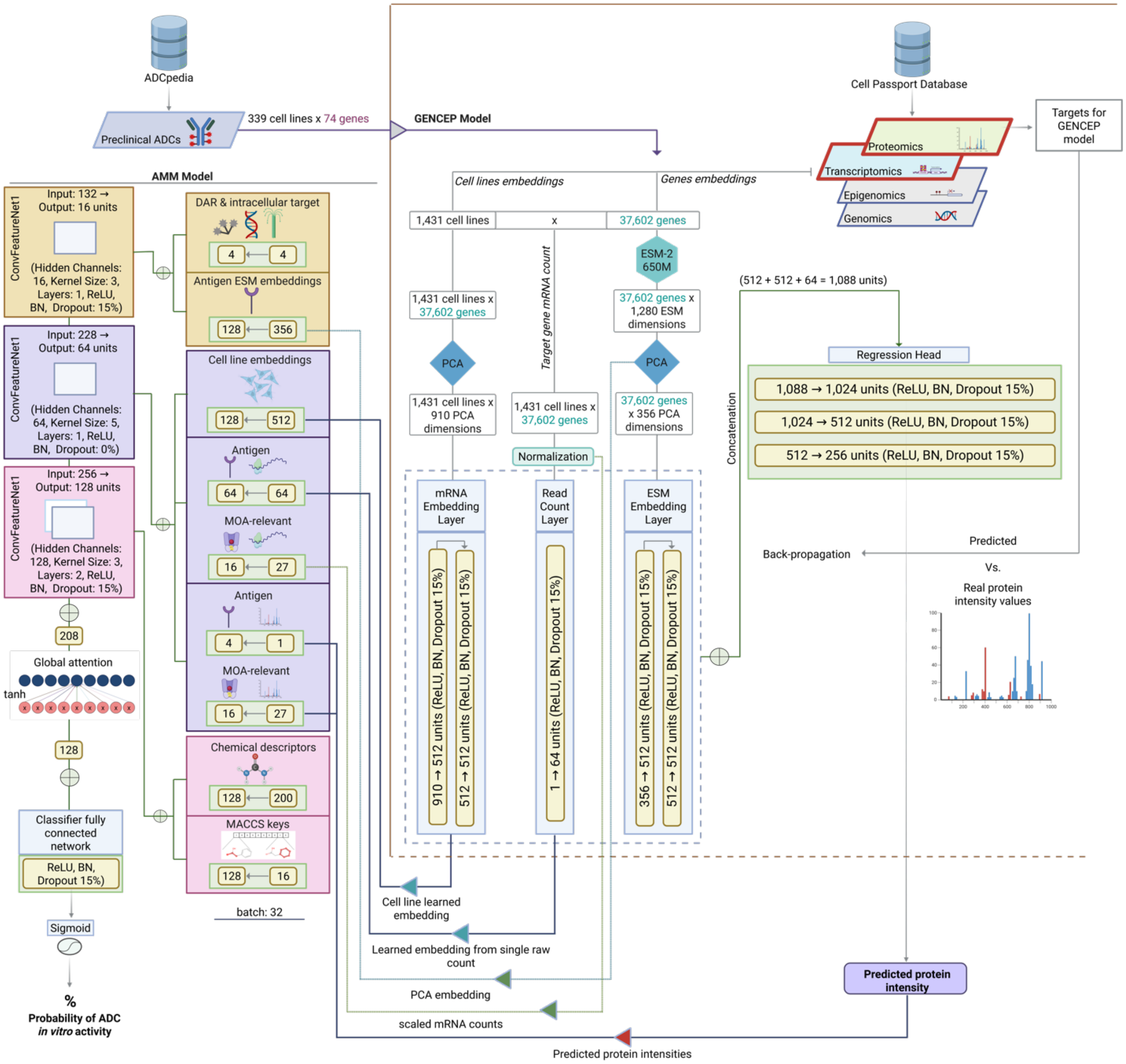

***Fig. S2. Schematic of GENCEP and AMM models architectures.***

*(**A**) Embeddings and outputs from GENCEP's expression predictions are fed into the AMM model, which processes these and other biological and physicochemical features via convolutional layers, to predict ADC activity as a classification task (≤10 nM = 1; >10 nM = 0). (**B**) The GENCEP model predicts protein expression intensities, leveraging mRNA counts, ESM embeddings, and target gene mRNA counts as inputs that are first processed by separated dense layers and then concatenated as an input for a regression head.*

### The evolution of ADC design and potency

Post-curation analysis of ADCpedia revealed significant trends in the evolution of linker-payload combinations over the ~25-year search span. Between 2001-2009, ADC research focused heavily on DNA damaging payloads like calicheamicin paired with acetyl-butyrate (AcBut) linkers. These linkers exploit the elevated intracellular glutathione levels to reduce the incorporated disulfide bonds and enable payload release (*1*). ADCs using this combination, such as gemtuzumab-ozogamicin (GO), were associated with $EC_{50}$ values below 10 nM and were predominantly tested on leukemic cell lines (*e.g.*, HL-60). In contrast, anti-microtubule payloads were paired with more diverse linkers, exhibited higher $EC_{50}$ values, and were tested on solid tumor cell lines (*e.g.*, A549 and U251). This likely reflects early challenges in optimizing anti-microtubule ADCs compared to the more developed AcBut-calicheamicin linker-payloads at the time. This stage of ADC development aligns with GO being the first clinically approved ADC in 2000, albeit it was withdrawn in 2010 due to fatal toxicities and then re-approved in 2017. Early exploration of polyethylene glycol (PEG) incorporation into linker structures also emerged during this interval.

From 2010-2018, research shifted toward protease-cleavable linkers, particularly dipeptide-based systems. This period saw extensive use of PEG moieties likely addressing challenges in conjugating hydrophobic small molecules to mAb side chains with a potential aim at higher DARs. There was also significant exploration of linker conjugation strategies involving lysine residues, which likely followed the development and success of trastuzumab emtansine (T-DM1) (*2*). Anti-microtubule payloads paired with cysteine conjugation strategies also gained prominence during this period and coincides with the success of brentuximab vedotin (BV) (*3*). However, the ADCs during this period showed diverse $EC_{50}$ values and were tested across both solid and liquid tumors. This likely reflects a phase of experimentation to identify effective payloads and optimize their pairing with various linker technologies for different tumor types.

The period from 2019 and onward is marked by a notable effort to increase payload diversity, with the inclusion of newer generations of DNA damaging agents and microtubule inhibitors. The valine-citrulline dipeptide coupled to the self-immolative para-aminobenzyl spacer (vc-PAB) became the linker most present, likely due to its ability to enable payload release without residual linker and/or antibody components and its improved controlled release relative to hydrazone-based linkers (*4*). This period also was characterized by novel discovery research focused on refining payload-linker pairings to maximize ADC efficacy against traditionally difficult to treat solid tumors such as glioblastoma (*e.g.*, SNB75), metastatic lung cancer (*e.g.*, NCI-H1930), and colorectal cancer (*e.g.,* HT-29). However, the ADC $EC_{50}$ values were highly variable and reflects the ongoing challenges in optimizing not only ADC structural combinations, but also the emerging importance of unique tumor type interfaces.

Regarding cytotoxicity, the field initially attempted to pair antibodies with traditional chemotherapeutics with micromolar $EC_{50}$ value range like doxorubicin however these ADCs had limited clinical anti-tumor activity (*5*). The five records in ADCpedia for ADCs with doxorubicin have $EC_{50}$ values ranging from 800-1250 nM. As previously described, this prompted a shift to develop ADCs with ultra-potent $EC_{50}$ values in picomolar range in the early 2000s (Fig. S2A). The notable shift, based on publication dates, occurred around 2010. ADC *in vitro* activities appeared to settle between 0.1 nM and 10 nM. However, upon closer inspection, ADC $EC_{50}$ values varied >300-fold (Fig. S2B). This wide distribution highlights areas of confusion within the field on what constitutes a 'potent' ADC design for a given target antigen and tumor type and is further complicated due to inconsistent conditions in cytotoxicity assays, and as optimization remains empirical, there is a need for cost-effective systematic strategies for the field to move forward (*6, 7*).

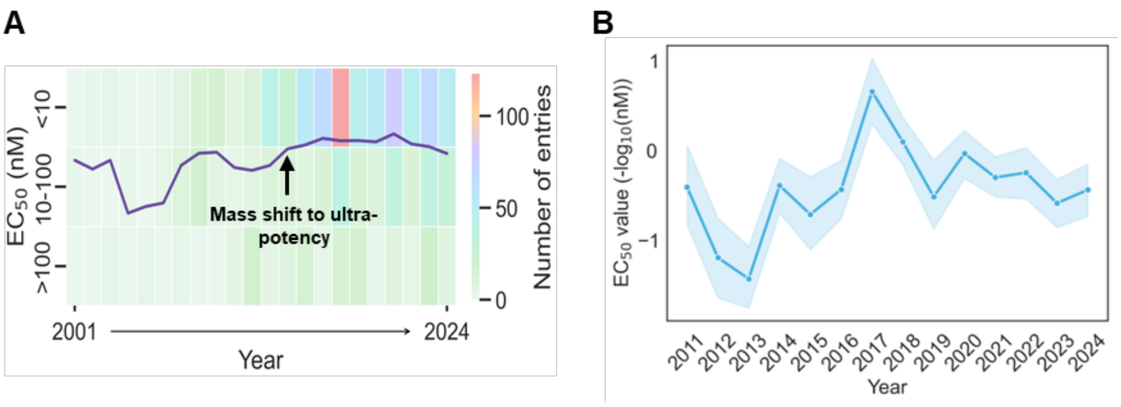

***Fig. S3. Historical ADC EC$_{50}$ values over time.***

*(**A**) Temporal trend of mean EC$_{50}$ values (line) for ADCs published from 2001-2024. Heat map showing the number of entries per year and stratified by <10 nM, 10-100 nM, and >100 nM EC$_{50}$ value bins. Arrow indicates the mass shift to ultrapotency. (B) Zoomed view of mean EC$_{50}$ values (-log$_{10}$(nM)) for ADCs from 2011-2024 indicated >300 variation.*

### The payload - ADC cytotoxicity relationship

Although DNA and microtubules are common intracellular targets for both ADCs and traditional chemotherapeutics, a key distinction lies in the high cytotoxicity of the payloads delivered by ADCs. These highly potent agents can overcome intrinsic resistance in certain tumor types to conventional chemotherapeutic mechanisms, which have been developed and optimized over decades for specific cancers. (*8*). Analyzing the tumor cell lines in ADCpedia revealed significant variability in EC$_{50}$ values across tumor types and highlights the challenge of matching specific payload structures to a given tumor context.

For ADCs delivering microtubule inhibitors, there were several tumor types that were sensitive with EC$_{50}$ values below 10 nM (Fig. S4A). DM1 is a derivative of maytansine, whose *in vitro* activity is 100- and up to 270-times more potent than traditional chemotherapeutic microtubule inhibitors vinca alkaloids and paclitaxel, respectively (*9-11*). Monomethyl auristatin E (MMAE) and monomethyl auristatin F are peptide analogs of dolastatin 10, which showed ultra-cytotoxic activities against human cancer cell lines (*12, 13*). As anticipated, breast and blood cancer cell lines were sensitive to microtubule inhibitor-incorporated ADCs with the majority of EC$_{50}$ values ≤10 nM. This aligns with the current indications for T-DM1, and Polatuzumab vedotin (Pola-V). There was considerable variation for many tumor types, possibly reflecting investigational ADCs that did not achieve the potency of more successful or approved counterparts. Notably, prostate cancer was the most sensitive tumor type, while sarcomas, mesothelial, kidney, and colorectal were notably less sensitive with ADC mean EC$_{50}$ values approaching 100 nM. Interestingly, lung cancers were distinctly divided in sensitivity between small cell and non-small cell types and explains the large variation in ADC cytotoxic potency. The mean $EC_{50}$ values for ADCs targeting non-small cell lung cancer were highly potent bordering just below 1 nM. In contrast, ADCs targeting small cell lung cancer cell lines displayed $EC_{50}$ values approaching 100 nM. Other cancers such as ovarian and oral cavity cancer cell lines had lower mean sensitivities compared to breast cancer and non-Hodgkin's lymphoma cell lines, albeit with significant variability, and suggest cancers can be targets for future ADC development. In contrast, colorectal tumor cell lines exhibited the least mean sensitivity, which aligns with colorectal cancer known as inherently resistant against anti-microtubule chemotherapy (*14*). However, the variability is wide and potentially indicates a microtubule inhibitor-incorporated ADC with a different design format (*e.g.,* linker) may prove otherwise.

For ADCs delivering DNA-damaging payloads, there were 13 tumor types that had been tested (Fig. S4B). It is currently thought the reason for why ADCs transporting calicheamicin payloads have had limited clinical effectiveness against solid tumors is due to poor tumor penetration and/or accessing of the DNA at concentrations below dose-limiting toxicity (*15, 16*). GO is indicated for the treatment of adult patients with newly diagnosed or relapsed/refractory CD33-positive acute myeloid leukemia (AML), and in pediatric patients (*17*). In general, DNA-damaging-incorporated ADCs had a mean $EC_{50}$ value approaching 1 nM for hematologic malignancy cell lines and support the effectiveness in constructing ADCs against AML. However, the blood cancer subtype, acute lymphoblastic leukemia, displayed much less sensitivity against these ADCs. Melanoma was highly resistant against DNA-damaging-incorporated ADCs, suggesting that DNA-damaging-based ADCs may not be ideal for this tumor type and that the cytotoxic potency itself and not the systemic dosing issue, should be considered. Nevertheless, such profound contrasts reinforce that ADCs delivering DNA damaging agents are under explored.

Topoisomerase I inhibitors represent a unique class of agents in the ADC landscape, with potencies that test the definition of 'ultra-toxic' (*18*). Topoisomerase I was originally validated as a cancer target when tumor cells died when treated with camptothecin, which was limited by its poor solubility and unacceptable toxicity (*19*). Both payloads SN38 and Dxd for the currently approved sacituzumab govitecan (SG) and trastuzumab deruxtecan (T-DXd), respectively, are camptothecin derivatives. SN38 is the active component of irinotecan and, importantly, is listed in the mid-to-high nM range and distant from the sub-nM payloads targeting DNA and tubulin (*18*). Comparatively, DXd, which is a derivative from exatecan, has been reported to be 10-fold more potent that SN38 (*20, 21*). Unsurprisingly, topoisomerase I inhibitor derivatives have been greatly pursued where at least 72 novel ADCs transporting more than 15 different camptothecin-based payloads, in combination with more than 21 different linkers across 24 different targets have been developed (*22*). Recently, several novel topoisomerase I inhibitors were reported all based on the camptothecin backbone and interestingly, exhibited sub-nM to low nM $EC_{50}$ values (*22*) and the available structures were integrated in ADCpedia.

An analysis of the ADCs delivering topoisomerase I inhibitors in ADCpedia revealed more uniform potencies with mean values all hovering around 10 nM (Fig. S4C). The number of tumor types of eight was notably less than the tumor systems evaluated with ADCs incorporated with microtubule inhibitors and DNA damagers. ADCs delivering topoisomerase I inhibitors were the most potent against thyroid cancer followed by hematologic malignancies and ovarian/endometrial/cervical-grouped cancers. There was wide $EC_{50}$ variability for lung cancers since, NSCLC tumor cell lines were sensitive while small-cell lung cancer cell lines were less sensitive. Interestingly, breast cancer was the most resistant tumor type with $EC_{50}$ values approaching 100 nM. The structural diversity of the topoisomerase I inhibitor-incorporated ADCs was significantly less compared to the ADCs incorporated with the two other payload types, as 129/138 ADCs were SG or T-Dxd or very similar (Fig. 2D). Additionally, these ADCs were almost all exclusively constructed with DARs of 8, eliminating DAR as a confounding variable and narrowing the focus to tumor biology matching. This further underscores the importance of identifying the optimal tumor type-payload combination when designing an ADC in early development.

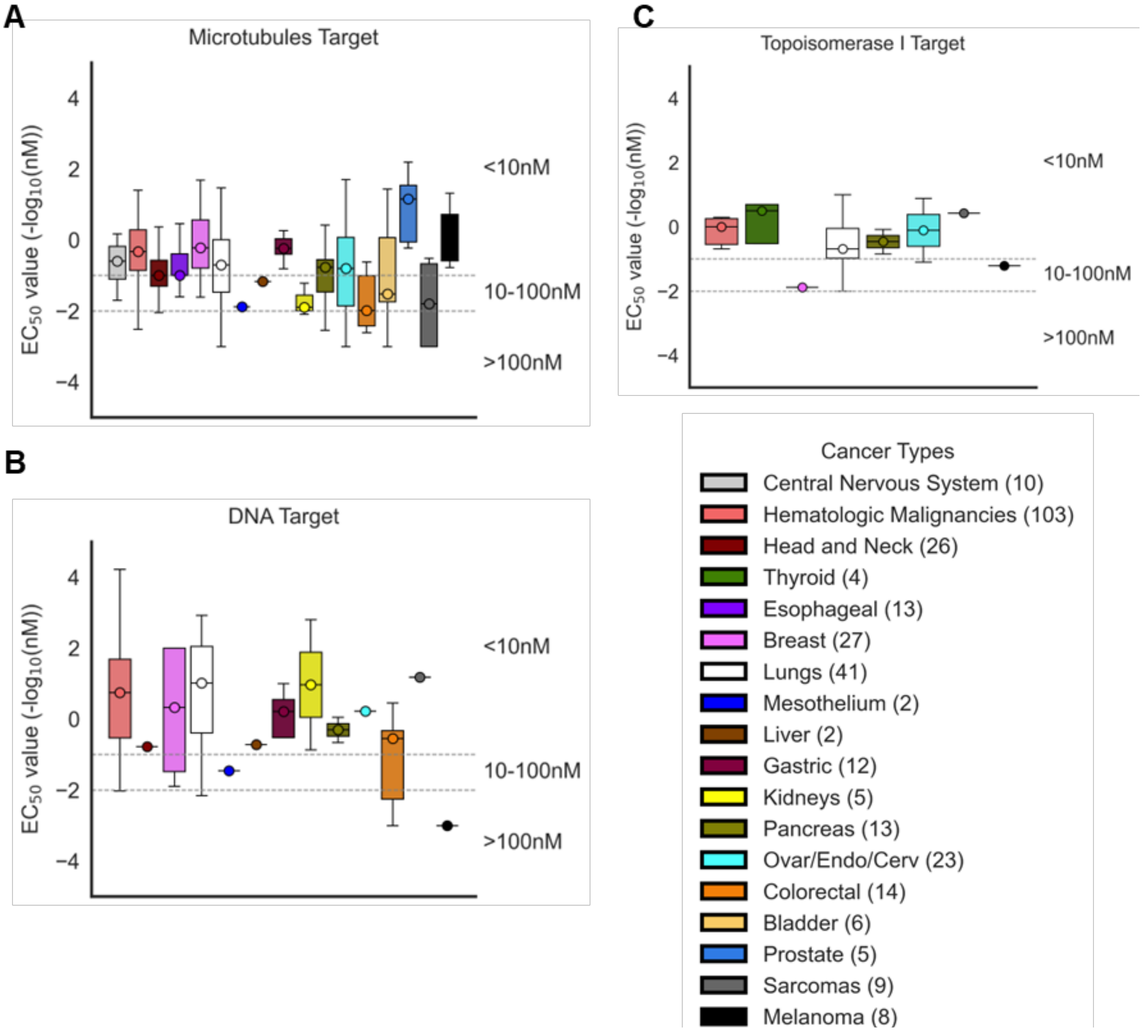

***Fig. S4. The relationship between ADC potency and payloads across tumor types.***

*The $EC_{50}$ values for ADCs in ADCpedia and the relationship with payload intracellular targets (**A**) microtubule inhibitors, (**B**) DNA damagers, and (**C**) topoisomerase I inhibitors. The zone between the dashed lines represents the 10-100 nM $EC_{50}$ value range. Above and below this zone are the $EC_{50}$ values <10 nM and >100 nM, respectively. Legend lists the tumor types by color.*

**The linker - ADC cytotoxicity relationship**

Linkers play a pivotal role in ADC design, substantially influencing both the stability and release of the cytotoxic payload. Although the past two decades ushered significant innovation in linker chemistry the types are overwhelmingly dominated by a few liner types ADCpedia such as mc-vc-PABC, SMCC, and CL2A (Fig. 2D).

There were 18 tumor types tested with cleavable linker-based ADCs (Fig. S5A), while only seven tumor types tested with non-cleavable linker-based ADCs (Fig. S5B). The ADCs incorporating cleavable linkers included those susceptible to cysteine-specific cathepsins (*i.e.*, valine-citrulline), cysteine- and serine-specific cathepsins (*e.g.*, GGFG), glutathione- (*i.e.*, AcBut) and prodrug-like (*e.g.*, hydrolysable CL2A) payload release mechanisms. As a group, these types of ADCs varied greatly in cytotoxic potency across all tumor types. The mean $EC_{50}$ values indicated that prostate cancer, B-cell lymphomas, breast cancer, NSCLC, AML, and small cell lung cancer cell lines are highly sensitive, albeit with wide variation. In contrast, ovarian and colorectal cancers exhibited notably less sensitivity, also with wide variability. Although mesothelial cancer displayed less sensitivity, there were only two ADC records.

The large $EC_{50}$ value differences may be immediately due to the diversification of the cleavable linkers contained in ADCpedia. For instance, novel amino acid combinations and sequence lengths linkers such as the GGFG linker used in T-Dxd (*21*), which is sensitive to both cysteine and serine proteases, it is not known how efficient cleavage and payload release is compared to other cleavable linkers. On the other hand, non-cleavable linkers require efficient lysosome ADC delivery and digestion to liberate the payload. For non-cleavable linker-incorporated ADCs, all seven tumor types displayed sensitivities of ≤10 nM. Therefore, in ADCpedia, the overall comparison of non-cleavable and cleavable linkers revealed a striking variability in cytotoxic potency and evolving linker chemistries, which was more notable with cleavable linker-based ADCs, most likely reflective of the different release mechanisms for these linkers.

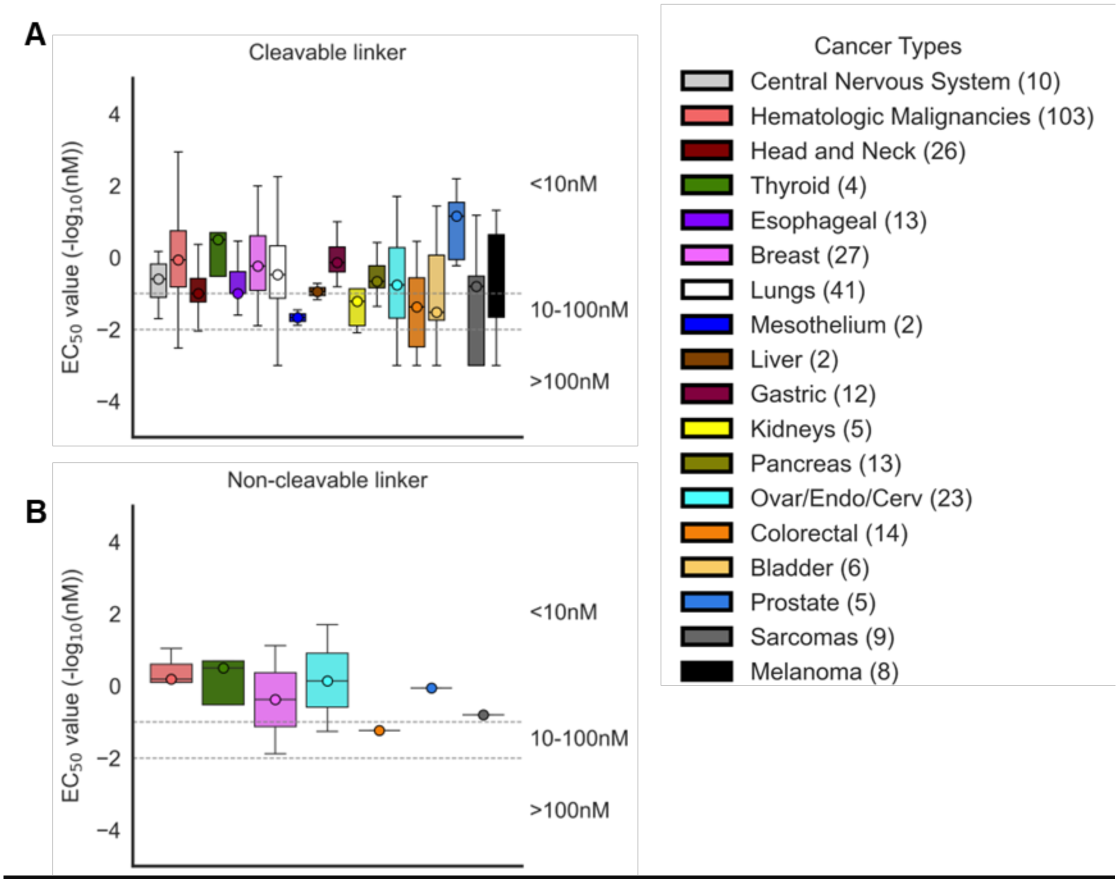

***Fig. S5. The relationship between ADC potency and linker types across tumor types.***
*The $EC_{50}$ values for ADCs in ADCpedia and the relationship with payload (**A**) cleavable (**B**) non-cleavable linkers. The zone between the dashed lines represents the 10-100 nM $EC_{50}$ value range. Above and below this zone are the $EC_{50}$ values <10 nM and >100 nM, respectively. Legend lists the tumor types by color.*

**<u>The DAR - ADC cytotoxicity relationship</u>**

DAR is greatly influential to ADC design as it effects overall potency, stability, and pharmacokinetics. The DARs in ADCpedia ranged from 0.8-9.0, with a mean DAR of 3.73. The most frequent DARs were 2.0, 3.4, 4.0, and 8.0 (Fig. S6). Importantly, both ultra-toxic and moderate-toxic payloads have been developed across this DAR spectrum over the past two decades. Interestingly, when DAR values were analyzed across tumor types and target antigens, no consistent patterns emerged. For cytotoxicity patterns, ADCs with DARs of 2 and 4 were more frequent with both high potency ($EC_{50}$ <1 nM) and low potency ($EC_{50}$ >100 nM). This suggest that the DAR at the 2-4 range did not uniformly correlate with enhanced cytotoxic potency.

Importantly, higher DARs were not associated with increased cytotoxic potency. This lack of association can be partially attributed to the inherent differences in free payload potency. For example, highly potent payloads such as PBD dimers achieve strong cytotoxic effects even at very low DARs of 1 (*23, 24*). In contrast, less potent payloads like SN38 are almost exclusively assembled with DARs of 8, which is thought necessary to compensate for reduced potency (*18*). Because ADCs with DARs of 8, particularly with ADCs delivering topoisomerase I inhibitors, are more recent advancements and are underreported in the current ADC literature, they are not as represented to potentially reveal emerging trends in this specific type of ADC design, especially at lower DARs.

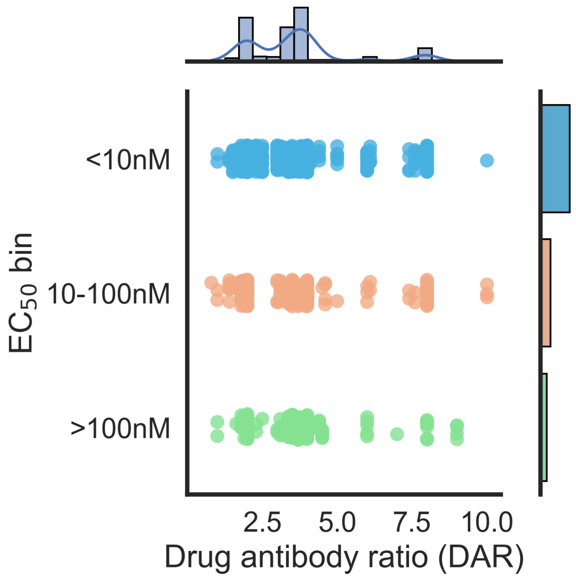

***Fig. S6. ADC cytotoxic potency variability as a function of DAR.***

*$EC_{50}$ values are binned across different DAR values. The top x-axis describes the distribution of DAR values across ADCpedia. The right y-axis represents the distribution of ADCs in ADCpedia.*

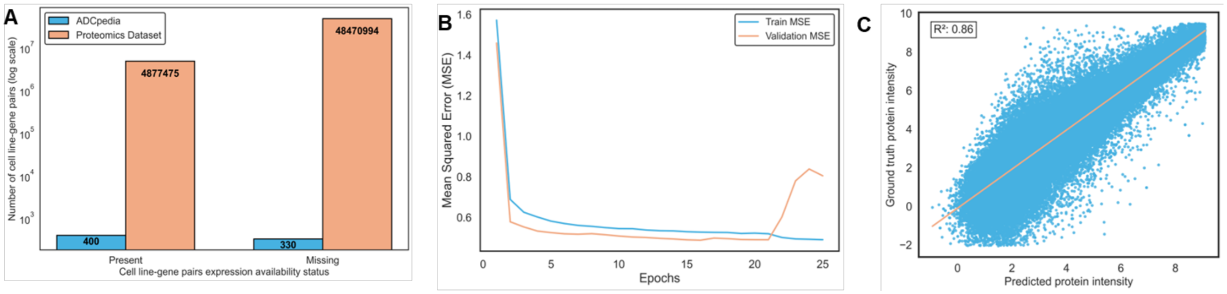

***Fig. S7. Expansion of proteomics data with GENCEP.***

*(A) Bar plot illustrating the total number of cell line-gene pairs with missing and present protein intensity values in the Cell Model Passport (blue = ADCpedia antigens and peach = overall). (B) Hyperparameter optimization and training over 24 epochs with early stopping. Learning curve of GENCEP on training (blue) and validation (peach) sets. (C) GENCEP-predicted intensities versus ground truth values on a set of ~400,000 random proteins from all cell lines in ADCpedia.*

**<u>The relationship between predicted target antigen intensity and ADC cytotoxicity</u>**

**HER2**

HER2 served as a benchmark to standardize the abundance of the antigens listed in ADCpedia. HER2 is a model antigen due to the amplification of the HER2/neu oncogene and the resulting high cell surface antigen densities, which have been directly correlated to outcomes for patients treated with trastuzumab (*25, 26*). Moreover, the relationship between ADC cytotoxicity in various levels of HER2 expressing tumor cells has previously been studied (*27*). For example, the HCC1937 and MCF-7 breast cancer (*28, 29*) cell lines are often accepted as 'low/negative' HER2-expressing cells used in studying T-DM1 (*30, 31*) and T-DXd (*27*).

To demonstrate the accuracy of the GENCEP-predicted intensities with the available ground truth intensities, the HER2 real/predicted intensities were 2.96/2.77 for MCF-7 and 3.24/3.16 for HCC1937 breast cancer cell lines. For the ovarian cancer cell line SK-OV-3, which has been used as a HER2 'high'-expressing model (*27*), the real/predicted intensities were 8.34/7.74. Therefore, standardized metrics for predicted intensities were created into high (≥7.5), mid (≥2.78-7.4), and low/negative (<2.78) expression levels across all tumor cell lines (Fig. S8A).

Notably, HER2 intensity stratification revealed that fewer cell lines than anticipated were predicted to express high levels with reported ADC activities. Tumor cell lines NCI-N87 (gastric carcinoma) and UACC-812 (breast ductal carcinoma) expressed very high HER2 levels, above 10 (Fig. S8A). At the mid expression level most HER2-positive cell lines belonged to breast cancer types. When evaluating the mean HER2 intensities for different cancer systems, novel insights became apparent. The average HER2 intensity was the highest at 3.94 for breast cancer, which was lower than the 4.24 mean intensity for all ADC-challenged tumor cell lines in ADCpedia (Fig. S8A and S8B). This indicated that other tumor types existed with attractive HER2 expression profiles. For example, blood and gastric cancers were potential attractive targets as their HER2 predicted intensities were 3.56 and 3.3, respectively. Additionally, lung, renal, melanoma, central nervous system (CNS), and female reproductive system-based cancers had mean HER2 intensities of ~3.0. Additional tumor types except for esophageal, pancreas, colorectal, and prostate cancers were all potential tumor types for the development of effective ADCs as their mean HER2 intensities were ≥2.78 (Fig. S8B). Interestingly, these means were all lower than the global mean of the tumor cell lines that had been empirically tested with investigational ADCs. This indicates that ADC development strategies could be biased against high-expressing HER2-positive tumor cell lines and not reflective of the overall expression within specific tumor types. Therefore, considering that there are scant records of ADC activities in many of these tumor types, there is considerable space for future anti-HER2 ADC development.

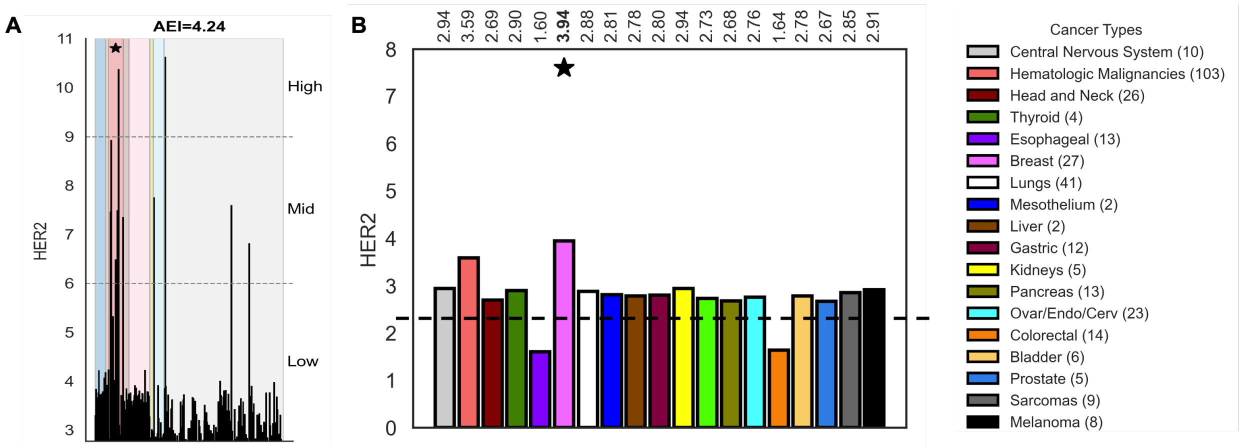

***Fig. S8. Predicted HER2 intensities across tumor cell lines.***

*(**A**) Diversity of the GENCEP-predicted intensities for HER2 in tumor cell lines challenged by an ADC in ADCpedia. (**B**) The averaged GENCEP-predicted intensities for each organ system. The number of cell lines is listed in Figure S1. Star indicates the tumor indication for the approved ADCs T-DM1 and T-DXd. AEI = Average expression intensity.*

For HER2-targeted cases, while ADCs tested against low HER2-expressing cell lines consistently exhibited $EC_{50}$ values >100 nM, the insightful revelation was that no clear distinction was observed between mid and high expression levels and their associated potencies (Fig. S9A). This demonstrates that even for a well-studied antigen like HER2, relationships between expression at a targetable intensity and efficacy remain ambiguous.

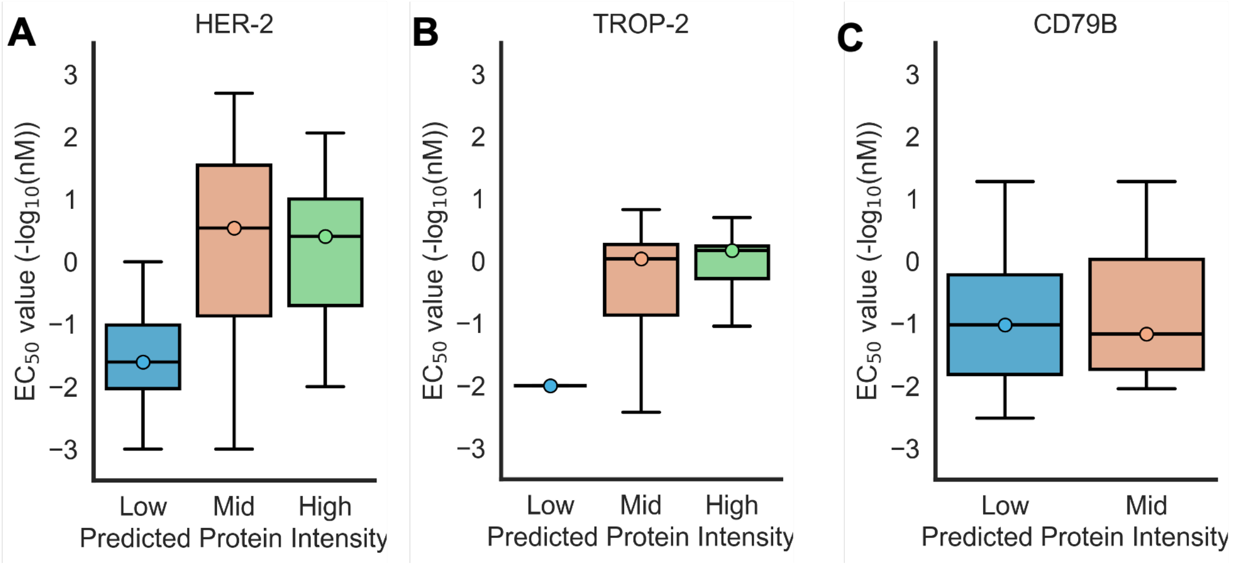

***Fig. S9. Relationship between GENCEP-predicted antigen intensity and $EC_{50}$ values in ADCpedia.***

*Boxplots evaluating the association between available $EC_{50}$ values (-log(EC50)) in ADCpedia across (A) HER2, (B) CD79b, and (C) TROP2 across their intensity tiers.*

### TROP2

Beyond HER2, GENCEP determined that the target antigens for approved ADCs all have intensities in the mid-to-low range and represents a significant under explored space for ADC design. For example, Trop2 is a 46-kDa monomeric glycoprotein whose expression has been reported in ~30 different tumor types, using different and semi-quantitative methods (*32*), and is the target of the approved ADC SG. SG was approved in 2021 for adult patients with Trop2-positive advanced and/or metastatic urothelial and triple-negative breast cancers based on the results from the TROPHY-U-01 (*33*) and ASCENT (*34*) studies, respectively. Unfortunately, SG was withdrawn for urothelial cancer as it did not meet the primary endpoints in the post-approval Phase 3, TROPiCS-04 study. Interestingly, there was no requirement nor measure of Trop2 for the TROPHY-U-01 study (*33*). However, archival tumor samples from patients enrolled in the TROPHY-U-01 study revealed that patient responses did not depend on Trop-2 expression levels, based on immunohistochemistry (*35*). Specifically, there was no statistical significance between groups based on high, mid, and low Trop2 expression and objective response rates, progression-free survival, and overall survival. Strikingly, patients with tumors with high Trop2 expression had poorer survival outcomes. A recent study simulating SG pharmacokinetics suggest that the instability of the CL2A linker in SG contributes to premature release of SN38, leading to excessive systemic exposure and severe adverse events (*36*) and detailed below in Section 14; cytotoxicity assay standardization. Additionally, when expressed at high levels in ovarian cancer, Trop2 inhibits apoptosis by increasing the expression of Bcl-2 and decreasing the expression of Bax (*37*), suggesting that absolute highest expression may be the most difficult tumor cells to kill. Therefore, Trop2 targeted ADCs warrant alternative designs that may be more effective by interconnecting with more underlying biological parameters.

In ADCpedia, many tumor cell lines overexpressing Trop2 and that had been challenged by an ADC reached the high expression level threshold (Fig. S10A). Several more tumor cell lines fell within the mid-level expression range. Notably, the mean predicted Trop2 intensity for ADC-challenged tumor cell lines was 5.50, higher than HER2. Extending to cell lines with no recorded ADC challenges, all organ systems and tumor types except for esophageal and colorectal cancers had mean Trop2 intensities well above the 2.77 cut-off (Fig. S10B).

There was also a notable overlap with $EC_{50}$ values for ADCs tested on cells with low- and mid-level Trop2 expression (Fig. S9B). This further demonstrates the ambiguity between expression and efficacy yet also presents potential broader therapeutic opportunities for Trop-2-targeting ADCs.

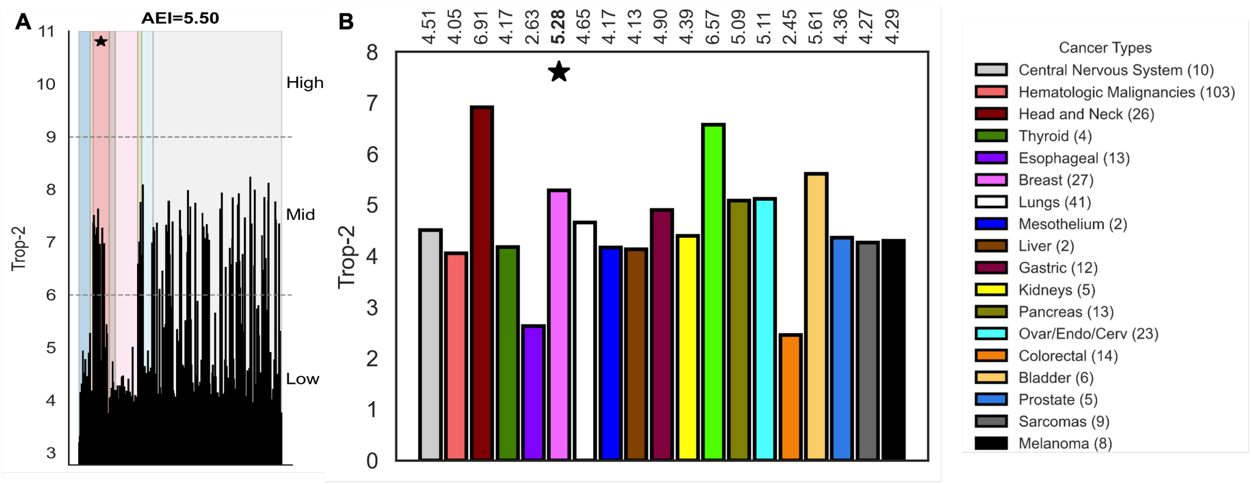

***Fig. S10. Predicted Trop2 intensities across tumor cell lines.***

*(**A**) Diversity of the GENCEP-predicted intensities for Trop2 in tumor cell lines challenged by an ADC in ADCpedia. (**B**) The averaged GENCEP-predicted intensities for each organ system. The number of cell lines is listed in Figure S1. Star indicates the tumor organ indication for the approved ADC SG. AEI = Average expression intensity.*

**CD79b**

CD79b is a component of the B-cell receptor complex and critical to the proper endocytosis of bound foreign antigens as part of the immune presentation pathway (*28*). CD79b is well documented to be exclusively expressed in immature and mature B cells but overexpressed in ≥80% of B cell-based neoplasms (*38, 39*). Pola-V is a clinically approved ADC specifically indicated for the treatment of patients with diffuse large B-cell lymphoma (DLBCL) based on results from the POLARIX trial (*40*). Preclinical testing of various Pola-V prototypes incorporating different combinations of linkers, payloads, and conjugation site on multiple CD79b-positive lymphoma cell lines revealed enhanced activity favored cleavable linker-incorporated ADCs (*41*). However, the CD79b expression levels on these cell lines were initially unknown. It was notable that a main reason for selecting the linker-payload design for Pola-V was that ADCs incorporating a non-cleavable linker exhibited poor internalization, and hence, ineffective payload release and anti-tumor activity (*42*).

It was subsequently reported that below a geometric mean fluorescence intensity threshold, CD79b-positive lymphoma cell lines were insensitive to anti-CD79b ADCs at a concentration of ~70 nM (*43*). Evaluating primary samples from patients with chronic lymphocytic leukemia, marginal zone lymphoma, hairy cell leukemia, follicular lymphoma, mantle cell lymphoma, and DLBCL showed that CD79b expression varied, was marginally higher than on normal B-cells, and there was a trend with relative higher expression correlated with more potent ADC $EC_{50}$ values (*43*). However, these expression levels were relative with absolute numbers missing.

In this work, GENCEP predictions revealed that the CD79b average expression intensity for hematologic malignancies was 3.28. Interestingly, all the tumor types apart from esophageal and colorectal cancers had considerably higher CD79b predicted intensities than the hematologic malignancies. However, there was no $EC_{50}$ value differences when stratifying cell lines into high and mid CD79b intensities (Fig. 9C). Taken together, CD79b is an attractive target for further ADC development for multiple types of cancer and, yet the relationship between expression and cytotoxic potency remains ambiguous.

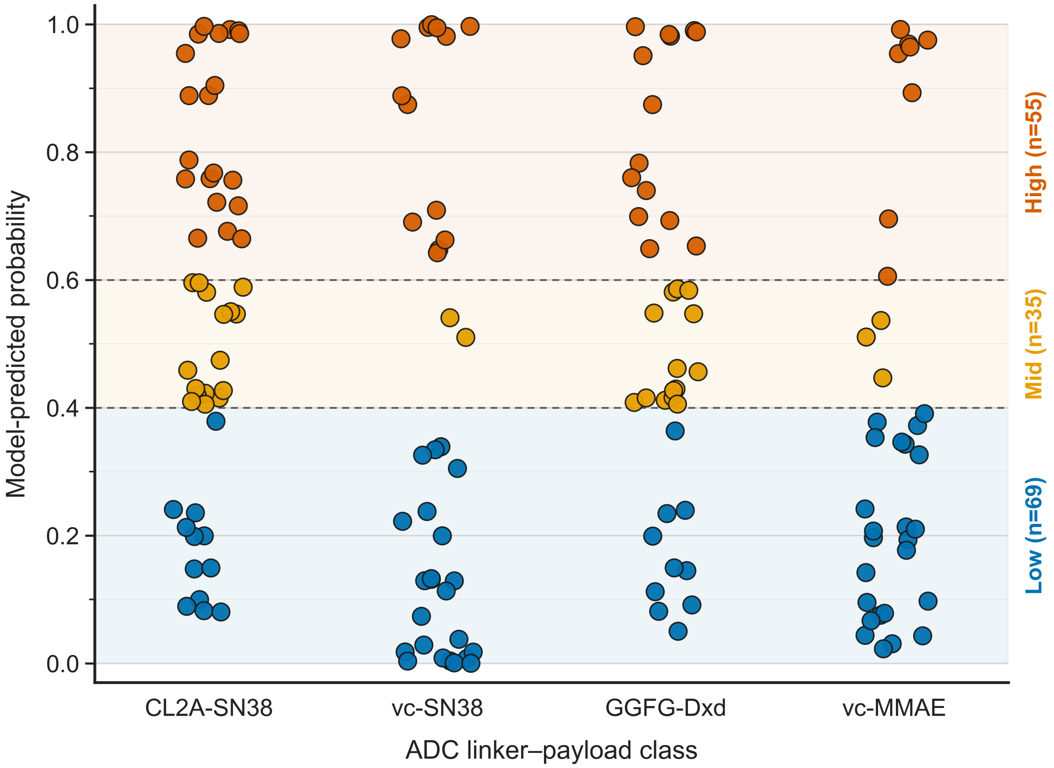

***Fig. S11. Probability distributions of payload-linker systems across all tested cell lines.***

*Each plotted point represents one ADC–cell line pair from the AMM model-predicted probability of activity likelihood outputs from 0.0 to 1.0. Each linker–payload class is in a distinct column and predicted-probability tiers (low (0.0 to 0.4), mid (0.4 to 0.6), or high (0.6 to 1.0)) are stratified by color. The number of points in each low, mid, and high tiers are in parentheses. High = likely sensitive, mid = intermediate, low = likely resistant.*

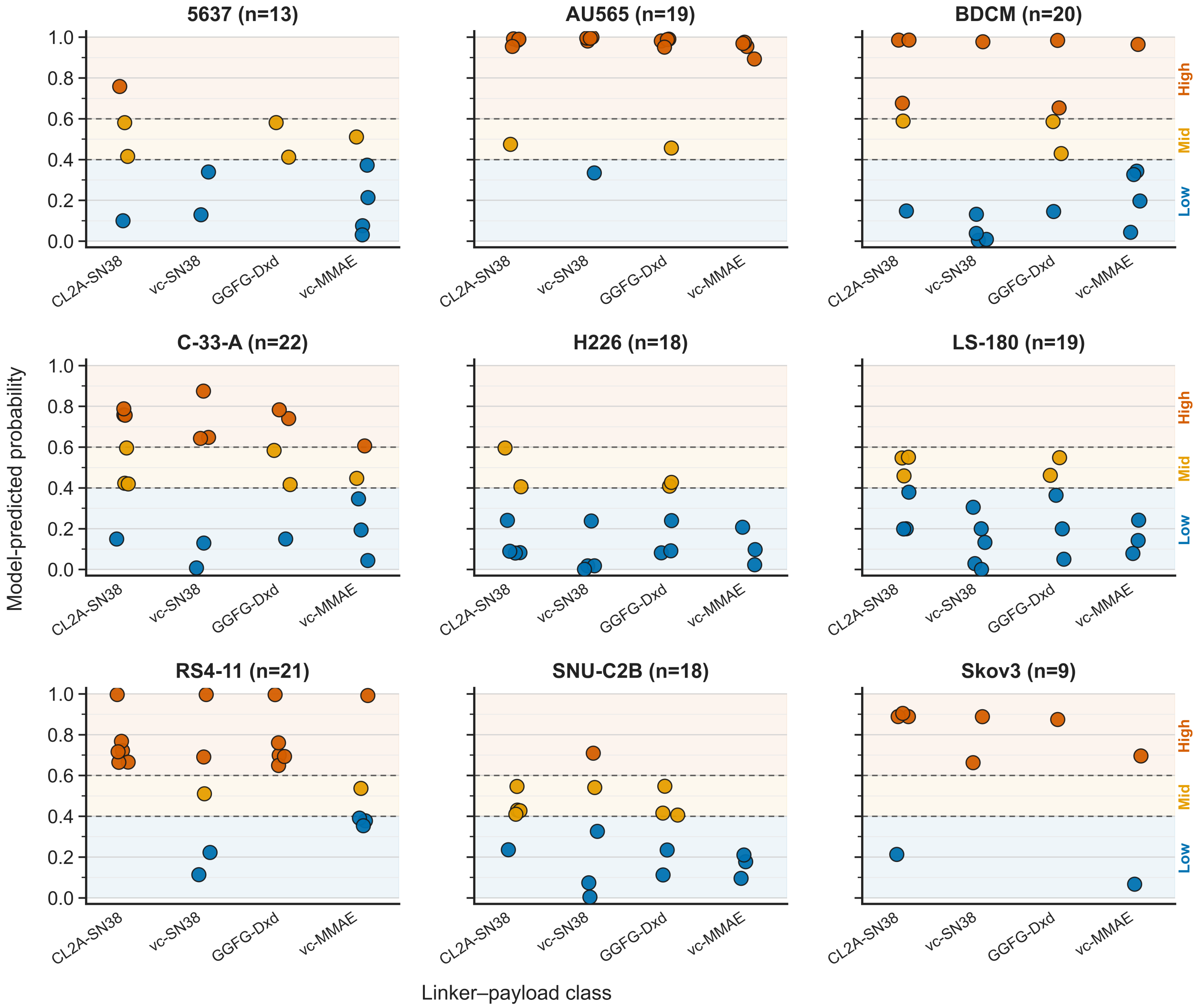

***Fig. S12. Probability distributions of payload-linker systems across each tested tumor cell line.*** *Each plotted point represents one ADC–cell line pair from the AMM model-predicted probability of activity likelihood outputs from 0.0 to 1.0. Each ADC is point stratified by linker–payload class is in a distinct column and predicted-probability tiers (low (0.0 to 0.4), mid (0.4 to 0.6), or high (0.6 to 1.0)) are stratified by color. The number of points in each low, mid, and high tiers are in parentheses. High = likely sensitive, mid = intermediate, low = likely resistant.*

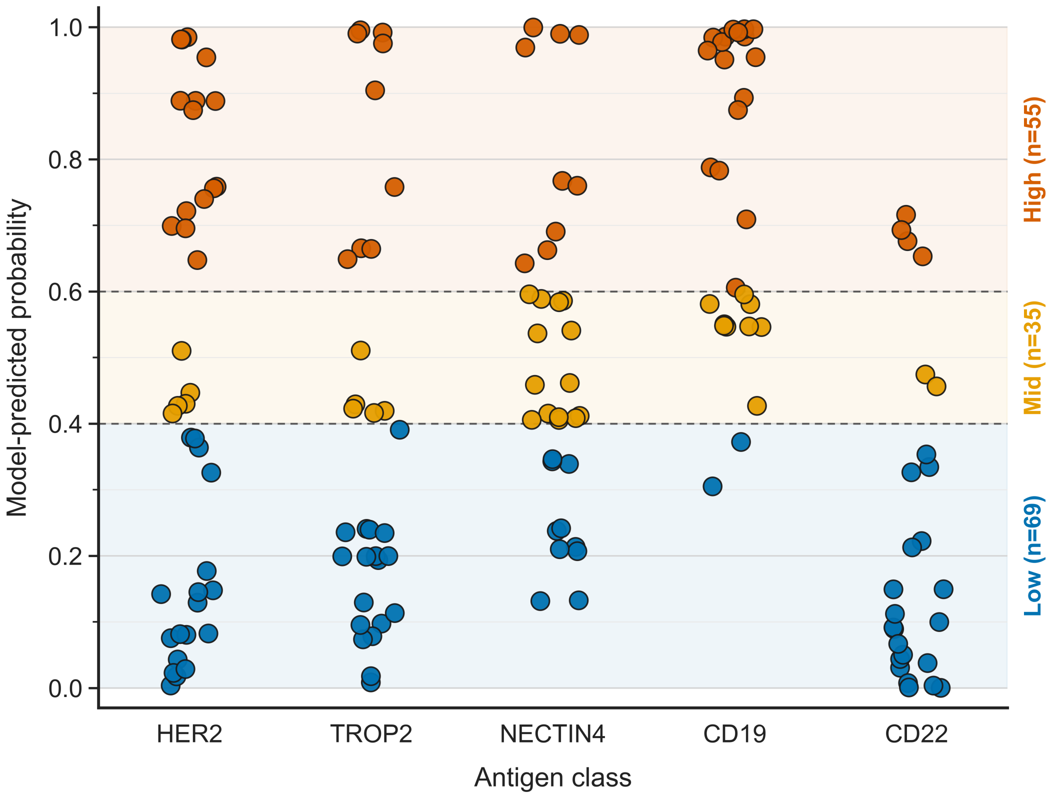

***Fig. S13. Probability distributions of designed ADCS, grouped by targeted antigen, across all tested cell lines.***

*Each point represents one ADC–cell line pair plotted by the AMM model-predicted probability of activity likelihood. Point shape denotes linker–payload class, and point color denotes predicted-probability tier: low (0.0 to 0.4), mid (0.4 to 0.6), or high (0.6 to 1.0). The number of points in each low, mid, and high tiers are in parentheses. High = likely sensitive, mid = intermediate, low = likely resistant.*

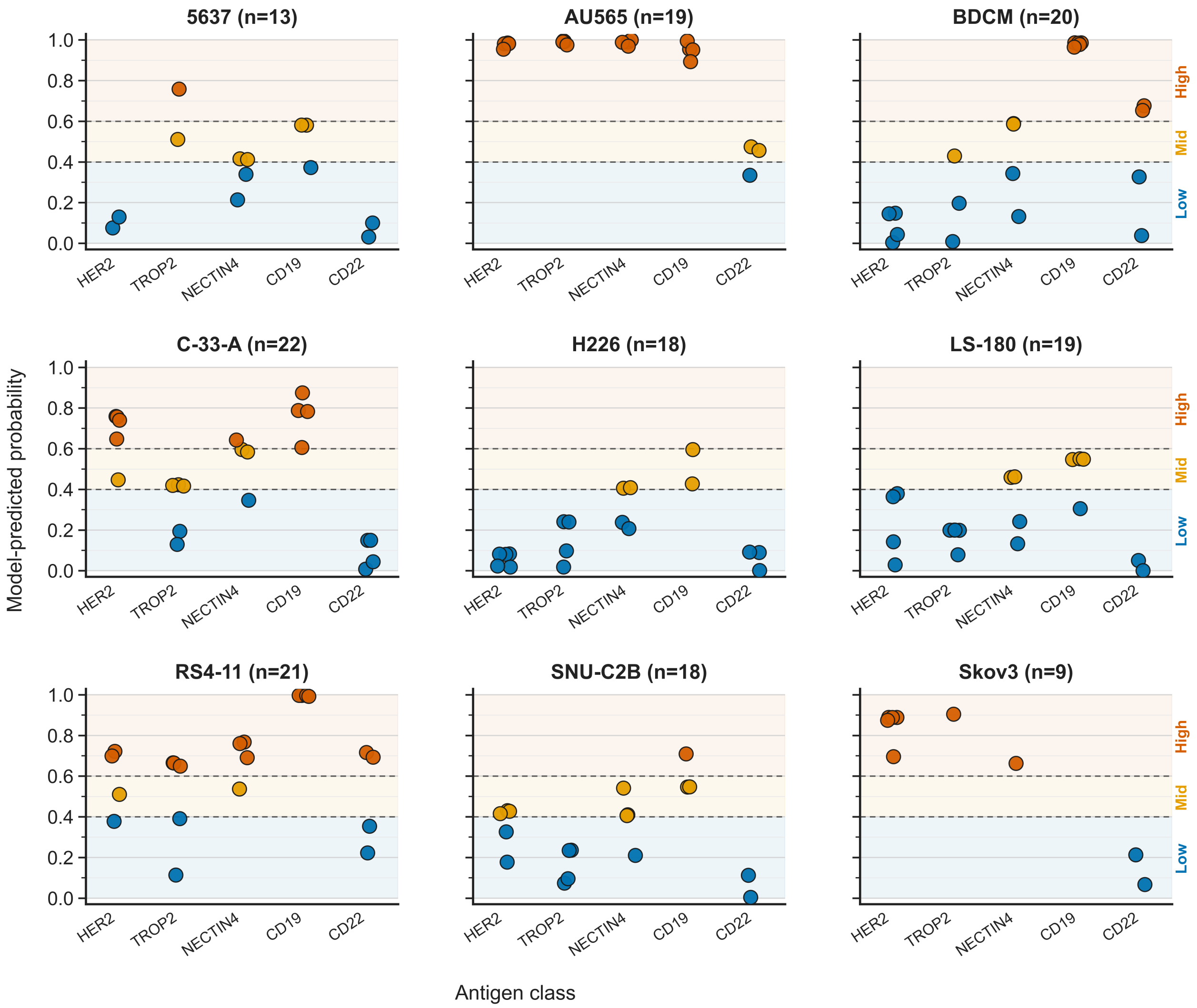

***Fig. S14. Probability distributions of designed ADCS, grouped by targeted antigen, within each tested tumor cell line.***

*Point color denotes predicted-probability tier: low (0.0 to 0.4), mid (0.4 to 0.6), or high (0.6 to 1.0). The number of points in each low, mid, and high tiers are in parentheses. High = likely sensitive, mid = intermediate, low = likely resistant.*

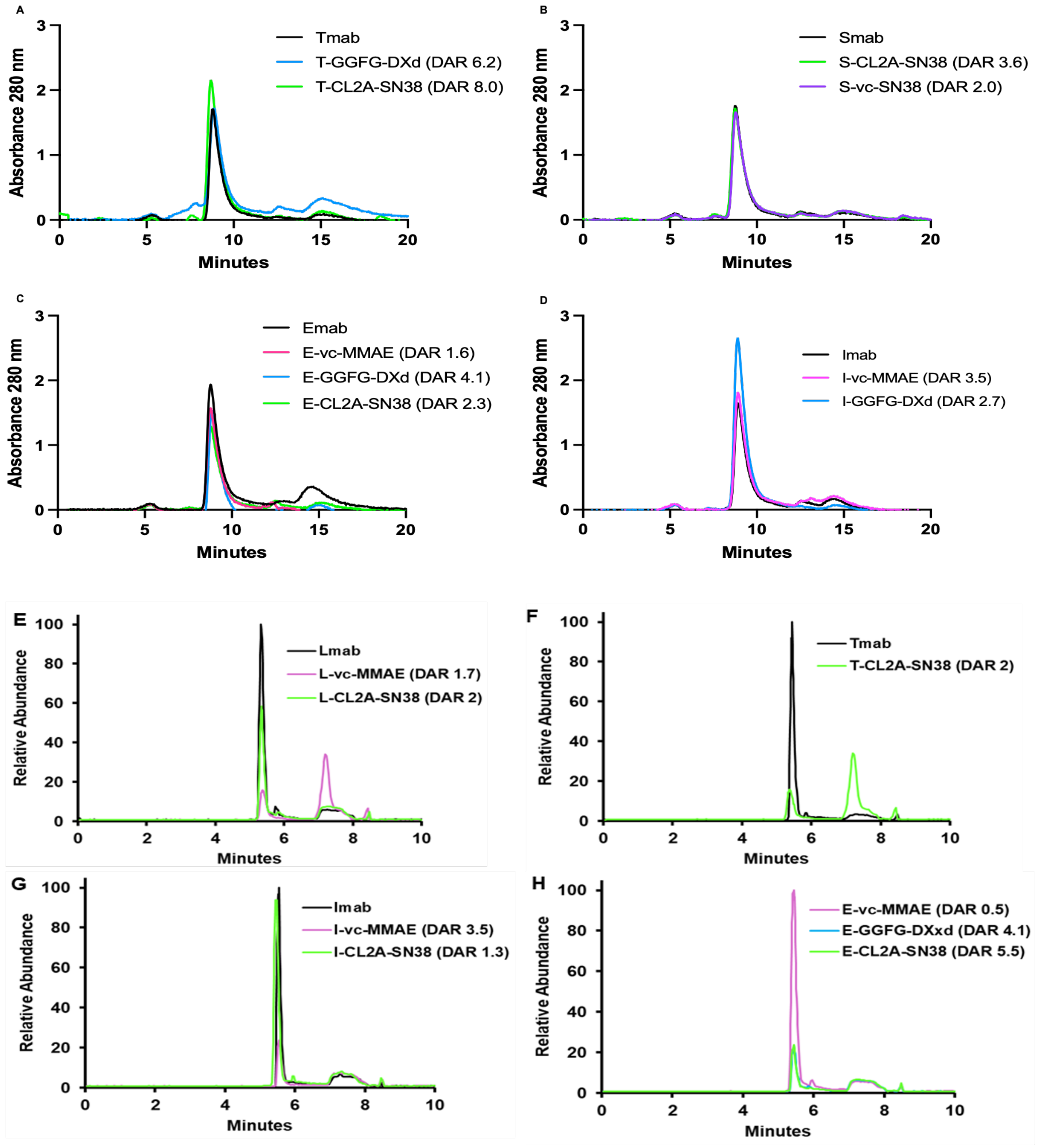

***Fig. S15. ADC uniformity profiles by SEC and nSEC-MS***

*SEC elution traces at 280 nm absorbance for A) trastuzumab (Tmab)-based ADCs, B) sacituzumab (Smab)-based ADCs, C) enfortumab (Emab)-based ADCs, and D) inotuzumab (Imab)-based ADCs. Native-SEC elution traces detected by mass spectrometry for E) loncastuximab (Lmab)-based ADCs, F) Tmab-based ADCs, G) Imab-based ADCs, and H) Emab-based ADCs.*

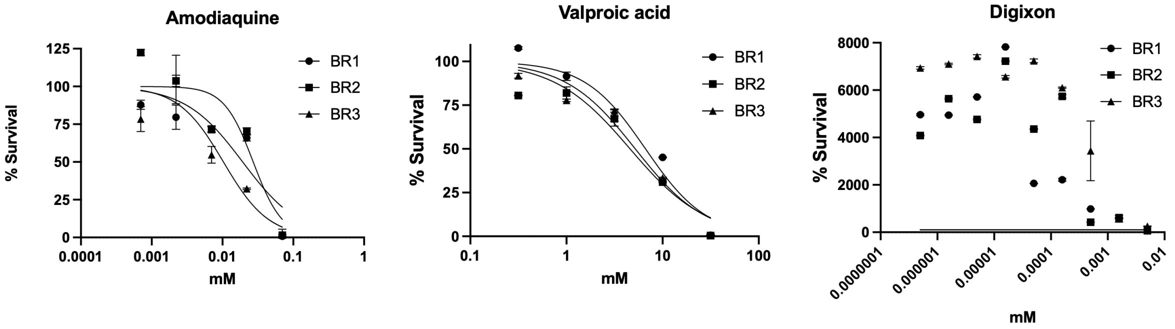

| Drug | $EC_{50}$ from 4pLL Prism | Published $EC_{50}$ |
|---|---|---|
| Amodiaquine | 0.019 ± 0.008 mM | 0.0185 mM (Proctor et al.) |
| Valproic acid | 5.478 ± 1.057 mM | 7.147 mM (Schürmeyer et al.) |
| Digixon | 0.047 ± 0.037 mM | <0.0002 mM (Proctor et al.) |

***Fig. S16. Cytotoxicity evaluation by Prism.***
*Inhibition curves and $EC_{50}$ values generated by non-linear regression of log-transformed drug concentrations (5-point [amodiaquine, valproic acid] or 9-point [digoxin] serial dilutions with top and bottom constrained to constant values of 100 and 0, respectively) from published literature and performed on our Prism software. BR = biological repeats.*

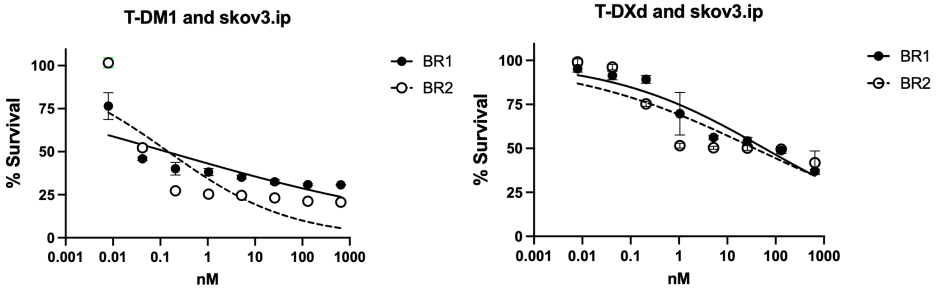

| ADC | $EC_{50}$ from 4pLL Prism |
|---|---|
| T-DM1 | 0.141 ± 0.009 nM |
| T-DXd | 47.7 ± 15.54 nM |

***Fig. S17. Cytotoxicity range evaluation with approved ADCs.***
*Inhibition curves and $EC_{50}$ values generated by non-linear regression of log-transformed drug concentrations on SKOV3.ip cells for T-DM1 and T-DXd. BR = Biological repeats.*

**ADC specificity evaluation.** T-vc-MMAE (DAR 2.2) demonstrated an $EC_{50}$ of 0.147 nM on SKOV3.ip cells with no calculable $EC_{50}$ on CHO-K1 cells, confirming antigen-dependent cytotoxicity consistent with the established proteolytic stability and receptor-mediated internalization mechanism of the vc-MMAE system (*3, 4*) (Fig. S19).

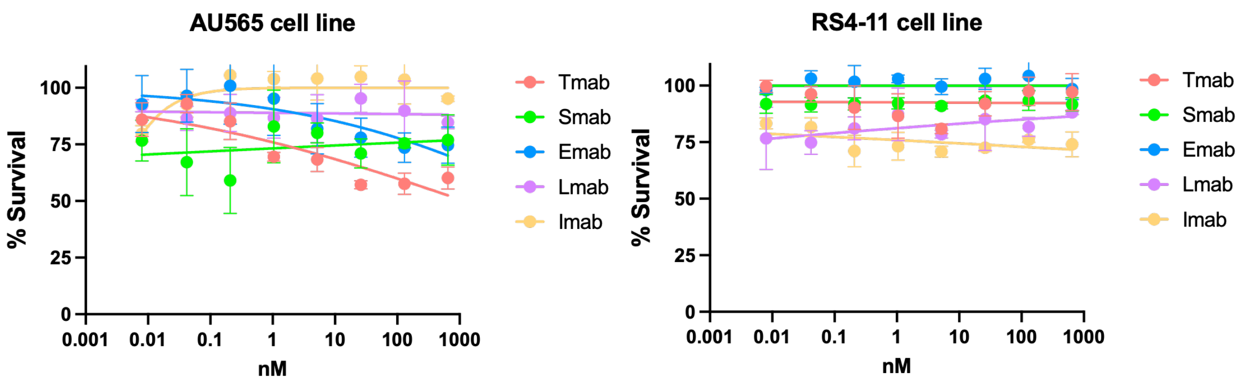

***Fig. S18. Cytotoxicity naked antibodies used for prospective validation.***
*Inhibition curves and $EC_{50}$ values generated by non-linear regression of log-transformed drug concentrations on sensitive predicted cell lines AU565 and RS4-11 for trastuzumab, sacituzumab, enfortumab, loncastuximab, and inotuzumab.*

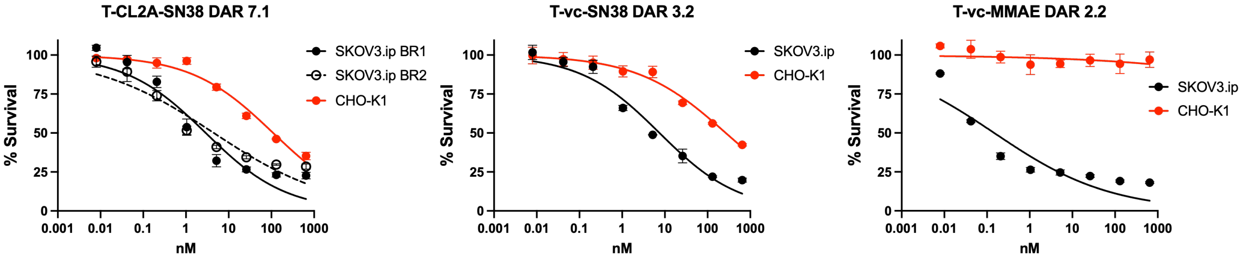

***Fig. S19. Cytotoxicity specificity of representative set of ADC designs for prospective validation.***
*Inhibition curves and $EC_{50}$ values generated by non-linear regression of log-transformed drug concentrations on SKOV3.ip and CHO-K1 cells for three ADC designs that would be used for prospective validation.*

However, the SN38-based constructs revealed a mechanistically informative cytotoxicity gradient. First, on SKOV3.ip cells across matched comparators: free SN38 (15.43 nM), free CL2A-SN38 linker-payload (28.81 nM), anti-CD20 rituximab-CL2A-SN38 (DAR 7.2; 57.13 nM), IgG-CL2A-SN38 (DAR 6.5; 118.1 nM), and T-CL2A-SN38 (DAR 7.1; 3.71 ± 1.12 nM) (Fig. S15 and S16). T-vc-SN38 (DAR 3.2) exhibited $EC_{50}$ values of 7.49 nM on SKOV3.ip and 267 nM on CHO-K1, yielding a 36-fold specificity margin. These results reflect the established prodrug pharmacology of SN38-based linker systems (*36, 44-48*). Specifically, a CL2A-SN38-based ADC released SN38 in PBS with a half-life of 8.6 h and a half-life of 10.8 h in human serum, revealing release spontaneous (*47*). Another study using sacituzumab-CL2A-SN38 reported that 50% of the drug was released every day (*44*). In cancer patients, the approved anti-Trop2 ADC, S-CL2A-SN38 had no correlation between efficacy and antigen expression and PK modeling has revealed that circulating levels of free antibody are 6-fold higher than intact ADC, where there stability of attached SN38 is approximately 14 h (*36, 49, 50*). Lastly, to the best of our knowledge there have not been a non-specific ADC controls reported for cytotoxicity. Therefore, the concordance between our results with free SN38, free CL2A-SN38, T-CL2A-SN38, R-CL2A-SN38, and IgG-CL2A-SN38 $EC_{50}$ values on SKOV3.ip and CHO-K1 cell lines confirms that cytotoxic activity reflects the composite mechanism of this linker-payload class, which is that spontaneous prodrug release is a major contributor to cytotoxicity and augments any antigen-mediated internalization contributions.

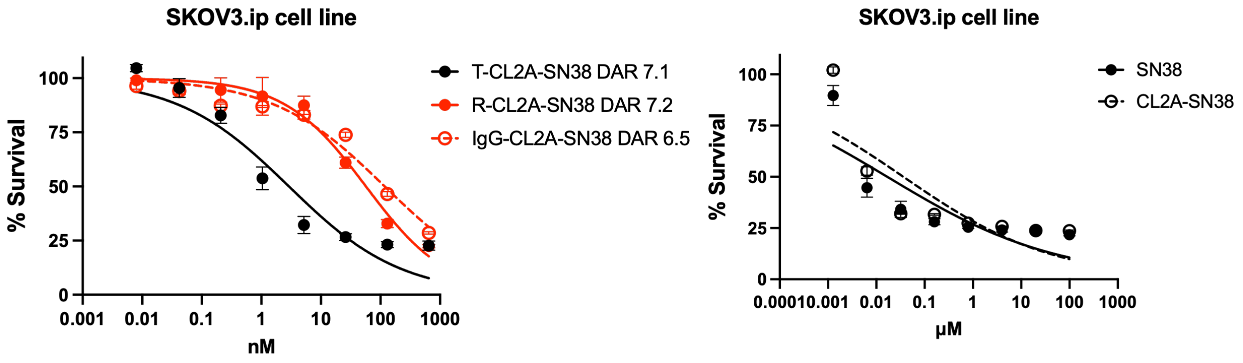

***Fig. S20. Cytotoxicity specificity on SKOV3.ip cells.***
*Left) Inhibition curves and* $EC_{50}$ *values generated by non-linear regression of log-transformed drug concentrations on SKOV3.ip cells for T-CL2A-SN38 and control ADCs, rituximab (R)-CL2A-SN38 and IgG-CL2A-SN38. Right) Percent survival of SKOV3.ip cells challenged with free SN38 or CL2A-SN38*

**SN38 release kinetics.** For the free CL2A-SN38 linker-payload at pH 4.0, SN38 release was rapid and near-complete, reaching 81.8% at 24 h and plateauing at 91.3% by 72 h, consistent with efficient acid-catabolyzed hydrolysis under conditions simulating the lysosomal environment (Fig. S21). For T-CL2A-SN38 at pH 4.0, release was progressive and substantial, reaching 48.7%, 58.9%, and 70.7% at 24, 48, and 72 h, respectively, reflecting the slower hydrolysis kinetics when the linker is conjugated to the antibody relative to the free compound.

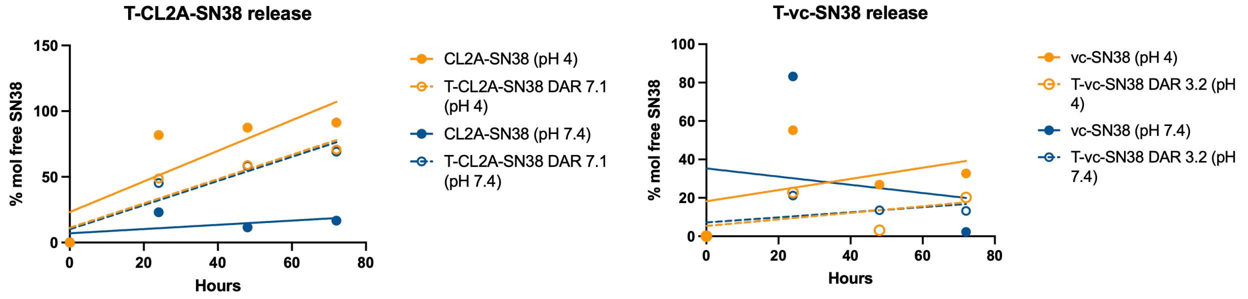

***Fig. S21. SN38 release kinetics.***
*The SN38 release kinetics for the CL2A-SN38 (left) and vc-SN38 (right) systems as linker-payload alone or when conjugated to trastuzumab in physiological PBS pH 7.4 (dark yellow) and pH 4.0 (blue).*

The most mechanistically significant finding was the release profile of T-CL2A-SN38 at physiological pH 7.4, which followed an essentially identical progressive trajectory to that observed at pH 4.0. The release was 45.3%, 58.3%, and 69.2% at 24, 48, and 72 h, respectively.

The near-equivalence of release at pH 7.4 and pH 4.0 indicates that SN38 dissociation from T-CL2A-SN38 proceeds through spontaneous hydrolysis that is largely pH-independent at these timescales, rather than requiring lysosomal acidification for efficient payload liberation.

These release kinetics directly account for the cytotoxic activity of non-binding antibody conjugates and the reduced antigen specificity margins observed for CL2A-SN38 constructs in the specificity evaluation. The concordance between free SN38 $EC_{50}$ values (15.43 nM), free CL2A-SN38 linker-payload $EC_{50}$ values (28.81 nM), and non-binding antibody-CL2A-SN38 $EC_{50}$ values (57.13 - 118.1 nM) on SKOV3.ip cells reflects the shared cytotoxic mechanism of spontaneously released free drug acting on cells regardless of antigen expression. The 15-fold potency enhancement of HER2-targeted T-CL2A-SN38 over non-binding rituximab-CL2A-SN38, and the 4-fold enhancement over free SN38, confirms that antigen-mediated internalization provides meaningful cytotoxic augmentation above the spontaneous release baseline, consistent with the prodrug model proposed for this linker class (*36*).

For T-vc-SN38, a more modest but detectable release of approximately 13% and 20% mol SN38 was observed at pH 7.4 and 4.0 over 72 h, consistent with the partial lability of the hydrozone attachment shared between vc-SN38 and CL2A-SN38 under physiological conditions and accounting for the 36-fold antigen specificity margin observed for this construct.

Collectively, the ADC characterization data present above are internally consistent and concordant with the published characterization of CL2A-SN38 and vc-SN38 ADC constructs produced by the same hydrazone-based conjugation chemistry and diafiltration purification strategy used here (*45, 48*). The spontaneous SN38 release kinetics, specificity margins, and free drug $EC_{50}$ concordance observed in this study fall within the ranges reported across the seminal CL2A-SN38 literature, confirming that the constructs used in prospective cytotoxicity testing were of equivalent quality and pharmacological character to those described in establishing this linker-payload class. These data collectively support the conclusion that the cytotoxic activity measured across 159 prospective ADC-cell line combinations reflects the genuine biological response to each construct under standardized real-world assay conditions.

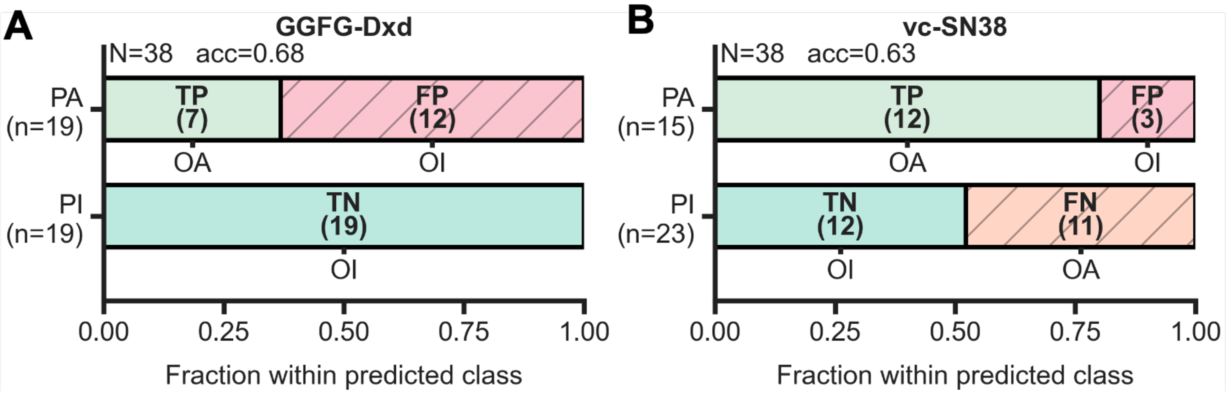

***Fig. S22. AMM prospective performances for GGFG-DXd- and vc-SN38-based ADCs.***
*Confusion matrix analyses for stratified performances for A) GGFG-DXd- and B) vc-SN38-based ADCs.*

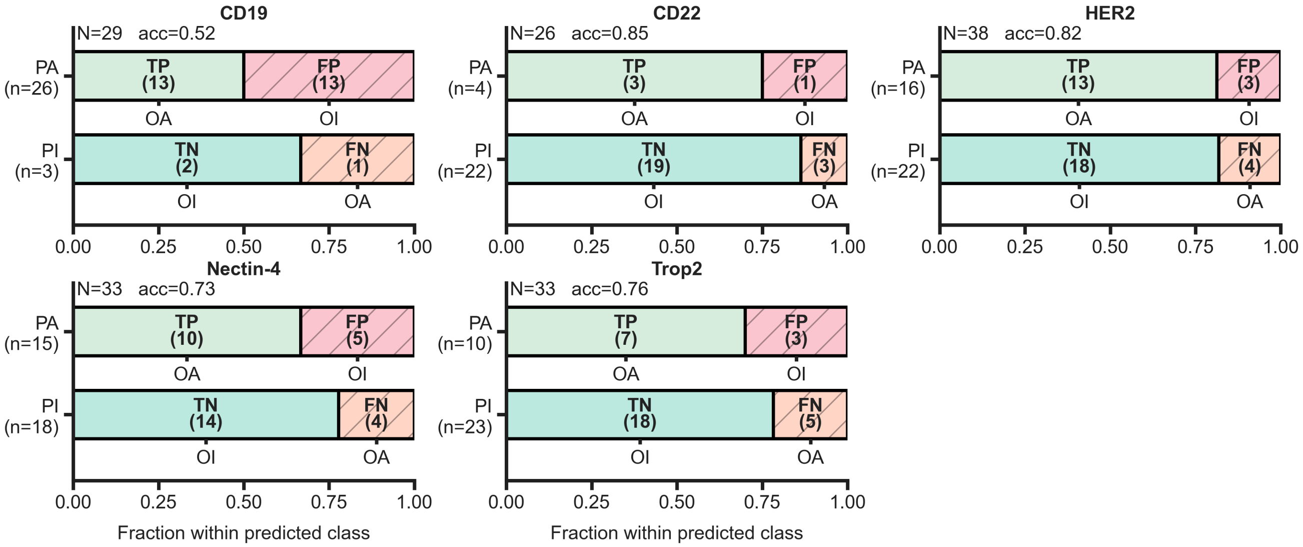

***Fig. S23. AMM prospective performances stratified by targeted antigens.***
*Confusion matrix analyses for stratified performances.*

### 15. Online interface

Allocation and deployment with Digital Research Alliance of Canada involved using a virtual machine allocation for public access, facilitated by a "Floating IP." A subdomain (server.adcpedia.com) was linked to this IP, and the application was developed with WordPress for the frontend and Django for the backend, integrating prediction models and data pipelines. Nginx and Gunicorn were used for deployment, while SSH managed the server. Project files were transferred, and settings were configured to handle cookies, CSRF, and CORS. Security rules for HTTP and HTTPS enabled requests on ports 80 and 443. Nginx was set up to redirect traffic to Gunicorn, serving the Django application. Static files were collected, database migrations executed, and SSL certificates obtained via Certbot for HTTPS security. File permissions ensured proper access, and multiple Gunicorn instances were able to maintain performance under heavy loads. Monitoring of Nginx logs helped detect and resolve issues. CSRF tokens secured POST requests, while CORS was configured to manage requests from different origins, essential for the Digital Research Alliance of Canada deployment.

### 16. Statistical analysis

Multivariable analytical methods were used to generate associations between ADC components and antigen expressions and $EC_{50}$ values. Continuous variables were compared using Pearson correlation coefficients and linear regression models, with model performance quantified by the coefficient of determination ($R^2$). For instance, correlations between scaled mRNA read counts and true protein intensities were assessed by scatter plots overlaid with regression lines, while differences in antigen expression across predefined $EC_{50}$ bins were examined using boxplots and strip plots. Outliers were identified and excluded based on Z-score and interquartile range criteria with the box encompassing the 75th interquartile range (IQR) and the mean indicated by horizontal lines in the boxes. Box whiskers span the 25th IQR. The $IC_{50}$ values were log-transformed ($pEC_{50} = -\log_{10}[EC_{50}]$) when appropriate. In addition, classification models for drug sensitivity (using an $EC_{50}$ threshold of 10 nM) were evaluated by ROC-AUC analyses for validation, test, blind, prospective datasets. Confusion matrices were further constructed for overall performance for generalizability, across cell group tiers, linker-payload systems, and certain cell line–specific analyses, generate statistical significance defined as $p<0.05$, sensitivity, specificity, positive predictive value, negative, predictive value, and likelihood ratio. Data was analyzed using Graphpad Prism and/or Python (pandas, SciPy, scikit-learn, and seaborn) for figure presentation. Concept figures were generated using BioRender.

### SUPPLEMENTARY TABLES

**Table S1.**
Best hyperparameters for the AMM model at 10 nM threshold, identified with Optuna hyperparameter optimization.

| Feature Stream | Data Inputs and Descriptors | Initial Linear Transform (Input → Output Dimensions) | CNN Architecture | CNN Output Dimensions |
|---|---|---|---|---|
| Stream 1 (DAR and ESM) | DAR and intracellular targets (4 features) | 4 → 4 for DAR and Targets | One-layer CNN with 16 hidden channels, kernel size of 3, dropout of 15 percent, followed by global average pooling | 16 |
| | PCA-reduced ESM antigen embeddings (356 features) | 356 → 128 for ESM | | |
| | | Both outputs concatenated into a single 132-dimensional input | | |
| Stream 2 (Antigen and MOA-Relevant Proteins) | Predicted antigen intensity (1 feature) | 1 → 4 for Antigen Intensity | One-layer CNN with 64 hidden channels, kernel size of 5, no dropout, followed by | 64 |

| | | | | |
|---|---|---|---|---|
| | | | global average pooling | |
| | mRNA cell line embeddings from the GENCEP model (512 features) | 512 → 128 for mRNA Cell Line Embeddings | | |
| | Antigen-specific mRNA read counts' embeddings from the GENCEP model (64 features) | 64 → 64 for Gene-Specific Read Counts | | |
| | Additional pharmacokinetic-related genes of mRNA and proteomic features from the GENCEP model (27 features) | 27 → 16 for Scaled mRNA Features | | |
| | | All outputs concatenated into a single 212-dimensional input | | |
| Stream 3 (Chemistry) | 200 RDKit descriptors (example : Balaban J, Bertz CT, Chi indices, EState, TPSA, etc.) | 200 → 128 for RDKit-based descriptors | Two-layer CNN with 128 hidden channels, kernel size of 3, dropout of 15 percent, followed by global average pooling | 128 |
| | 167-bit MACCS molecular fingerprints | 167 → 128 for Molecular Fingerprints | | |
| | | Both outputs concatenated into a single 256-dimensional input | | |
| Attention Mechanism | Produces a combined embedding across all streams | Attention layer with a hidden dimension of 128 | | Attention-weighted combined embedding |
| Fully Connected Classifier | Refines the combined embedding | Combined embedding → 128; Fully connected layer | Outputs binary probability for $EC_{50}$ less than 10 nM | |

| | | | |
|---|---|---|---|
| | | with dropout of 15 percent, ReLU activation, batch normalization, and sigmoid output | |
| Training Hyperparameters | Learning rate: 4.71E -4 | | Optimizer: Adam, Batch size: 32, Weight decay: 1E -5 |

**Table S2.**
Confusion matrix results for *in silico* internal test set.

| **Metric** | **Value** | **95% Confidence interval** |
|---|---|---|
| p-value | 0.0004 | Significant |
| Sensitivity | 0.667 | 0.454 – 0.828 |
| Specificity | 0.833 | 0.681 – 0.921 |
| PPV | 0.70 | 0.481 – 0.855 |
| NPV | 0.811 | 0.658 – 0.905 |
| LR | 4.0 | |

PPV = positive predictive value; NPV = negative predictive value, LR = likelihood ratio

**Table S3.**
Excel sheet (Table S3_retrospective validation.xlsx)

**Table S4.**
Confusion matrix results for external blinded retrospective test set.

| Metric | Value | 95% Confidence interval |
|---|---|---|
| p-value | 0.0009 | Significant |
| Sensitivity | 0.929 | 0.685 – 0.987 |
| Specificity | 0.857 | 0.487 – 0.974 |
| PPV | 0.929 | 0.685 – 0.987 |
| NPV | 0.857 | 0.487 – 0.974 |
| LR | 6.50 | |

**Table S5.**
Cell line information for prospective validation.

| Cell line (tumor type) | Sex, age | Media | ATCC link |
|---|---|---|---|
| AU565 (breast) | F, 43 | RPMI-1640 | https://www.atcc.org/products/crl-2351 |
| LS-180 (colorectal) | F, 58 | EMEM | https://www.atcc.org/products/cl-187 |
| SNU-C2B (colorectal) | F, 43 | RPMI-1640 | https://www.atcc.org/products/ccl-250 |
| SKOV3 (ovarian) | F, 64 | McCoy's | https://www.atcc.org/products/htb-77 |
| RS4;11 (acute lymphoblastic leukemia) | F, 32 | RPMI-1640 | https://www.atcc.org/products/crl-1873 |
| C-33-A (cervical) | F, 66 | EMEM | https://www.atcc.org/products/htb-31 |
| NCI-H226 (mesothelioma) | M, unspecified | RPMI-1640 | https://www.atcc.org/products/crl-5826 |
| BDCM (acute myeloid leukemia) | M, unspecified | RPMI-1640 | https://www.atcc.org/products/crl-2740 |
| 5637 (bladder) | M, 68 | RPMI-1640 | https://www.atcc.org/products/htb-9 |

**Table S6.**
Probability vs $EC_{50}$ comparison including the number of replicates performed for each ADC-cell line combination. Green = Correct prediction. Red = Incorrect prediction. ND = cytotoxicity experiment was not performed.

| ADC | 5637 | AU565 | BDCM | C-33-A | H226 | LS180 | RS4-11 | SKOV3 | SNUC2B |
|---|---|---|---|---|---|---|---|---|---|
| **T-CL2A-SN38** | 0 | 2 | 2 | 5 | 2 | 1 | 3 | 2 | 2 |
| **Probability** | | 0.9848 | 0.1477 | 0.7585 | 0.0824 | 0.3789 | 0.7216 | 0.8884 | 0.4300 |
| **EC$_{50}$ (nM)** | NP | 1.99 ± 0.94 | 0.63 ± 0.58 | 2.53 ± 0.98 /0.6367 | 12.3/35.7 | 13.58 | 1.13 ± 1.18 | 21.69/6.993 | 47.27/67.49 |
| **T-vc-SN38** | 1 | 3 | 4 | 3 | 4 | 3 | 4 | 1 | 4 |
| **Probability** | 0.1291 | 0.9812 | 0.0037 | 0.6475 | 0.0176 | 0.0286 | 0.5101 | 0.8882 | 0.3258 |
| **EC$_{50}$** | 8.541 | 7.55 ± 10.34 | 8.08 ± 6.93 | 8.7 ± 0.84 | 47.2 ± 24.1 | 3.81 ± 2.05 | 1.68 ± 0.6 | 1.036 | >100 |
| **T-GGFG-DXd** | 0 | 1 | 2 | 4 | 2 | 2 | 3 | 1 | 3 |
| **Probability** | | 0.9815 | 0.1450 | 0.7399 | 0.0814 | 0.3637 | 0.6991 | 0.8744 | 0.4153 |
| **EC$_{50}$** | NP | 1.99 ± 0.94 | >700 | >300 | >500 | 74.5 ± 4 | >100 | 2.143 | >1000 |
| **T-vc-MMAE** | 1 | 1 | 2 | 4 | 1 | 1 | 2 | 2 | 1 |
| **Probability** | 0.0753 | 0.9542 | 0.0431 | 0.4467 | 0.0228 | 0.1420 | 0.3775 | 0.6956 | 0.1770 |
| **EC$_{50}$** | 17.19 | 0.076 | >400 | 28.54 ± 18.04 | 519 | 21.11 | 339.4 ± 11.3 | 0.3 ± 0.22 | 788 |
| **S-CL2A-SN38** | 1 | 4 | 0 | 6 | 2 | 5 | 4 | 1 | 2 |
| **Probability** | 0.7581 | 0.9918 | | 0.4227 | 0.2408 | 0.1995 | 0.6654 | 0.9043 | 0.2356 |
| **EC$_{50}$** | 6.175 | 6.72 ± 4.66 | NP | 3.01 ± 2.3/11.54 ± 7.45 | 91.6 ± 7.2 | 8 ± 8.2/3.61 ± 2.19 | 2.86 ± 6.12/4.691 | 35.98 | >100 |

| | | | | | | | | | |
|---|---|---|---|---|---|---|---|---|---|
| **S-vc-SN38** | 0 | 2 | 5 | 4 | 3 | 1 | 3 | 0 | 3 |
| **Probability** | | 0.9952 | 0.0084 | 0.1293 | 0.0178 | 0.1996 | 0.1133 | | 0.0736 |
| **$EC_{50}$** | NP | 12.25 ± 7.56 | 7.4 ± 6.79 | 11.74 ± 3.91 | 39 ± 17.3 | 34.68 | 2.52 ± 1.03 | NP | >700 |
| **S-GGFG-DXd** | NP | 2 | 4 | 4 | 3 | 2 | 2 | 0 | 2 |
| **Probability** | | 0.9902 | 0.4292 | 0.4162 | 0.2395 | 0.1991 | 0.6489 | | 0.2344 |
| **$EC_{50}$** | NP | >400 | 44.03 ± 32.09 | 83.9 ± 21.11 | >500 | 273 ± 65 | 8.95 ± 1.99 | NP | >1000 |
| **S-vc-MMAE** | 1 | 3 | 3 | 2 | 2 | 1 | 2 | 0 | 1 |
| **Probability** | 0.5106 | 0.9753 | 0.1966 | 0.1935 | 0.0974 | 0.0783 | 0.3907 | | 0.0953 |
| **$EC_{50}$** | 0.03 | 2.61 ± 1.98 | >500 | >100 | 13.6 ± 0.5 | 557 | >1000 | NP | >1000 |
| **E-CL2A-SN38** | 1 | 3 | 2 | 7 | 3 | 3 | 3 | 0 | 3 |
| **Probability** | 0.4152 | 0.9899 | 0.5887 | 0.5957 | 0.4056 | 0.4587 | 0.7673 | | 0.4098 |
| **$EC_{50}$** | 15.71 | 2.95 ± 1.56 | 9.45 ± 1.1 | 5.66 ± 2.9 | 40.1 ± 17.6 | 1.93 ± 0.32 | 1.15 ± 0.83 | NP | >300 |
| **E-vc-SN38** | 2 | 3 | 5 | 3 | 1 | 1 | 3 | 2 | 3 |
| **Probability** | 0.3390 | 0.9994 | 0.1315 | 0.6426 | 0.2377 | 0.1326 | 0.6906 | 0.6625 | 0.5408 |
| **$EC_{50}$** | 9.99 ± 2.49 | 9.24 ± 14.42 | 2.95 ± 3.78 | 5.5 ± 4.23 | 53.41 | 2.92 | 4.24 ± 3.62 | 7.6 ± 2.17 | >200 |
| **E-GGFG-DXd** | 1 | 3 | 3 | 2 | 3 | 2 | 2 | 0 | 4 |
| **Probability** | 0.4117 | 0.9882 | 0.5859 | 0.5838 | 0.4083 | 0.4615 | 0.7600 | | 0.4057 |
| **$EC_{50}$** | 489.1 | >500 | 9.47 ± 3.24 | >100 | >700 | >200 | 37.02 ± 16.5 | NP | >1000 |

| | | | | | | | | | |
|---|---|---|---|---|---|---|---|---|---|
| **E-vc-MMAE** | 1 | 2 | 1 | 4 | 1 | 1 | 3 | 0 | 1 |
| **Probability** | 0.2134 | 0.9691 | 0.3429 | 0.3462 | 0.2070 | 0.2416 | 0.5365 | | 0.2100 |
| **$EC_{50}$** | 71.14 | 9.15 ± 0.01 | 422 | 399.03 ± 92.52 | >1000 | 904 | >1000 | NP | >1000 |
| **L-CL2A-SN38** | 1 | 1 | 5 | 4 | 2 | 7 | 4 | 0 | 2 |
| **Probability** | 0.5808 | 0.9545 | 0.9856 | 0.7877 | 0.5955 | 0.5466 | 0.9968 | | 0.5462 |
| **$EC_{50}$** | 7.081 | 128.7 | 3.61 ± 4.61/1.44 ± 1.48 | 7.01 ± 2.99 | 65.8 ± 34.2 | 5.82 ± 4.52/6.3 ± 3.96 | 1.02 ± 0.81 | NP | 154 ± 46 |
| **L-vc-SN38** | 0 | 1 | 2 | 4 | 0 | 1 | 1 | 0 | 2 |
| **Probability** | | 0.9947 | 0.9774 | 0.8745 | | 0.3050 | 0.9968 | | 0.7091 |
| **$EC_{50}$** | NP | 0.562 | 0.22 ± 0.09 | 6.72 ± 2.83 | NP | 2.71 | 0.49 | NP | 89.6 ± 79.3 |
| **L-GGFG-DXd** | 1 | 1 | 2 | 4 | 1 | 3 | 3 | 0 | 4 |
| **Probability** | 0.5811 | 0.9510 | 0.9842 | 0.7828 | 0.4269 | 0.5482 | 0.9963 | | 0.5471 |
| **$EC_{50}$** | 212.6 | 361.2 | 8.44 ± 3.07 | 332.3 ± 92.5 | >1000 | >400 | 7.31 ± 2.32 | NP | >400 |
| **L-vc-MMAE** | 1 | 2 | 1 | 5 | 0 | 0 | 1 | 0 | 0 |
| **Probability** | 0.3722 | 0.8930 | 0.9645 | 0.6057 | | | 0.9920 | | |
| **$EC_{50}$** | 70.18 | 526 ± 31.8 | 405.4 | 273.7 ± 91.9 | NP | NP | >1000 | NP | NP |
| **I-CL2A-SN38** | 1 | 2 | 3 | 6 | 2 | 0 | 2 | 1 | 0 |
| **Probability** | 0.0997 | 0.4743 | 0.6761 | 0.1492 | 0.0893 | | 0.7160 | 0.2127 | |
| **$EC_{50}$** | 3.273 | 11.35 ± 2.13 | 3.37 ± 1.67 | 15.76 ± 8.16 | >100 | NP | 3.63 ± 3.66 | 47.55 | NP |

| | | | | | | | | | |
|---|---|---|---|---|---|---|---|---|---|
| **I-vc-SN38** | 0 | 3 | 5 | 5 | 3 | 3 | 4 | 0 | 2 |
| **Probability** | | 0.3343 | 0.0377 | 0.0075 | 0.0008 | 0.0001 | 0.2223 | | 0.0036 |
| **$EC_{50}$** | NP | 259 ± 63.3 | 6.67 ± 3.16 | 15.03 ± 4.61 | 85.2 ± 55.7 | 10.8 ± 6.6 | 3.58 ± 4.56 | NP | 86 ± 29 |
| **I-GGFG-DXd** | 0 | 2 | 2 | 3 | 1 | 1 | 4 | 0 | 3 |
| **Probability** | | 0.4563 | 0.6531 | 0.1494 | 0.0914 | 0.0502 | 0.6929 | | 0.1121 |
| **$EC_{50}$** | NP | 184 | 61.4 ± 29.4 | >200 | >1000 | 220 | 9.07 ± 5.46 | NP | >800 |
| **I-vc-MMAE** | 1 | 0 | 3 | 5 | 0 | 0 | 1 | 1 | 0 |
| **Probability** | 0.0306 | | 0.3263 | 0.0438 | | | 0.3534 | 0.0667 | |
| **$EC_{50}$** | 23.15 | NP | >600 | 104.1 ± 29.8 | NP | NP | >1000 | 76.05 | NP |
